# Dynamic states of eIF6 and SDS variants modulate interactions with uL14 of the 60S ribosomal subunit

**DOI:** 10.1101/2022.07.29.502049

**Authors:** Jonah Elliff, Aparna Biswas, Sahiti Kuppa, Poonam Roshan, Angela Patterson, Jenna Mattice, Mathivanan Chinnaraj, Ryan Burd, Sarah E. Walker, Nicola Pozzi, Edwin Antony, Brian Bothner, Sofia Origanti

**Affiliations:** Department of Biological Sciences, Marquette University, Milwaukee, WI 53233, USA; Department of Immunology, The University of Iowa, Iowa City, IA 52242, USA; Department of Biology, Saint Louis University, St. Louis, MO 63103, USA; Department of Biochemistry and Molecular Biology, Saint Louis University School of Medicine, MO 63104, USA; Department of Chemistry and Biochemistry, Montana State University, Bozeman, MT 59717, USA; Department of Biological Sciences, State University of New York, Buffalo, NY 14260, USA

**Author notes:** Address correspondence to: Sofia S. Origanti, Department of Biology, Macelwane Hall 307, 3507 Laclede Ave., St. Louis, MO-63103. “The authors wish it to be known that, in their opinion, the first two authors should be regarded as joint First Authors”.

**Keywords:** eIF6/uL14/60S/Shwachman-Diamond, Syndrome/eIF6, somatic, mutations/Translation

## Abstract

Assembly of ribosomal subunits into active ribosomal complexes is integral to protein synthesis. Release of eIF6 from the 60S ribosomal subunit primes 60S to associate with the 40S subunit and engage in translation. The dynamics of eIF6 interaction with the uL14 (RPL23) interface of 60S and its regulation by somatic mutations acquired in Shwachman-Diamond Syndrome (SDS) is yet to be clearly understood. Here, by using a modified strategy to obtain high yields of recombinant human eIF6 we have uncovered the critical interface entailing 8 key residues in the C-tail of RPL23 that is essential for physical interactions between 60S and eIF6. Disruption of the complementary binding interface by conformational changes in eIF6 disease variants provide a mechanism for weakened interactions of variants with the 60S. Hydrogen-deuterium exchange mass spectrometry (HDX-MS) analyses uncovered dynamic configurational rearrangements in eIF6 induced by binding to RPL23 and exposed an allosteric interface regulated by the C-tail of eIF6. Disrupting a key residue in the eIF6-60S binding interface markedly limits proliferation of cancer cells, which highlights the significance of therapeutically targeting this interface. Establishing these key interfaces thus provide a therapeutic framework for targeting eIF6 in cancers and SDS.

## Introduction

Several *trans*-acting factors coordinate the assembly and maturation of the ribosomal subunits^1–4^. Eukaryotic translation initiation factor-6 (eIF6) is one such essential factor that is crucial for the biogenesis and maturation of 60S subunits^5–8^. It also functions as a 60S silencing or anti-association factor that sterically inhibits 60S association with the 40S subunit by disrupting inter-subunit bridges^9–14^. Release of eIF6 from 60S is thus critical to facilitate interactions with 40S and to permit the formation of translationally competent 80S monosomes^9–14^.

Impaired function of eIF6 contributes to the underlying pathology of certain cancers and inherited ribosomopathies: Shwachman-Diamond Syndrome (SDS) and a subset of pediatric T-cell acute lymphoblastic leukemia^15–18^. eIF6 is overexpressed in several cancers including colon, ovarian and breast cancers, and its enhanced expression is associated with a poor prognosis^19–26^. Partial loss of eIF6 limits the transformation efficiency of oncogenic H-Ras12V mutant and oncogenic Myc. A reduction in eIF6 levels in malignant pleural mesotheliomas and in Myc-induced lymphomas markedly impairs cell growth by inhibiting protein synthesis rates^20–22^. Since haploinsufficiency of eIF6 delays tumorigenesis without markedly affecting normal growth, it presents eIF6 as a viable therapeutic target with potentially minimal side effects^21^.

Thus, disrupting eIF6 activity has been proposed to be a therapeutic strategy for the treatment of certain cancers and SDS. This therapeutic strategy is especially significant for SDS that is predominantly driven by biallelic mutations in the Shwachman-Bodian-Diamond Syndrome (*SBDS*) factor^16, 18, 27^. SDS is an inherited disorder associated with bone marrow failure, exocrine pancreatic insufficiency, skeletal deformities, developmental defects, and a predisposition to myelodysplastic syndrome (MDS) and acute myeloid leukemia^28, 29^. Germline mutations in *SBDS* and *Elongation Factor like GTPase-1* (*EFL1*) hinder 60S maturation and prevent the release of eIF6 from nascent 60S leading to impaired subunit joining and reduced translational fitness^16–18, 28, 30^. It is likely that the release of eIF6 from recycled 60S subunits post-termination are also disrupted by SBDS deficiency^31^. Recent studies have detected somatic mutations in *EIF6* in the hematopoietic cells of SDS patients that either reduced eIF6 expression or weakened interactions of eIF6 with 60S^32, 33^. Acquisition of such *EIF6* mutations present a compensatory mechanism that rescue the ribosomal and translational defect of SBDS deficient cells^32, 33^. Thus, these studies highlight the importance of targeting eIF6 activity for a better prognosis for SDS patients either by genetic compensation or by therapeutic means.

A potent therapeutic strategy to disrupt eIF6 function is to block the critical interactions between eIF6 and 60S. However, to pursue such a targeted approach, there is a need to identify and experimentally interrogate the contributions of the residues in the binding pocket that are critical for eIF6 and 60S interactions. Structural studies show that eIF6 directly associates with RPL23 (uL14) and is proximal to the Sarcin-Ricin Loop (SRL) of the 28S rRNA and RPL24 (eL24) of 60S (Fig. 1a)^11, 34, 35^. One of the challenges towards performing extensive biophysical and biochemical characterization of the 60S-binding interactions is the inability to obtain sufficient quantities of full-length human eIF6^36^. Here, we describe a strategy to obtain milligram quantities of active full-length eIF6 by co-expression of the Trigger Factor (TF) chaperone. This development enabled us to probe eIF6 interactions with RPL23 through detailed biophysical characterization. Using surface plasmon resonance (SPR) and hydrogen-deuterium exchange mass spectrometry (HDX-MS) analyses, we have identified residues in eIF6 and in the C-terminus of RPL23 that are critical for eIF6-60S interactions. Our results show for the first time a dynamic transition state of eIF6 upon binding to RPL23 that entails conformational changes in regions that are disrupted by SDS mutations. In addition, circular dichroism (CD) analyses have uncovered the influence of the predominant N106S mutation and the key Tyr151 residue on the overall secondary structure of eIF6. Based on these analyses, we selectively targeted Tyr151 residue and show for the first time that perturbing a residue in the eIF6-60S interface markedly affects cancer cell proliferation rates. This further highlights the significance of targeting the interface as a therapeutic strategy.

**Figure 1.**
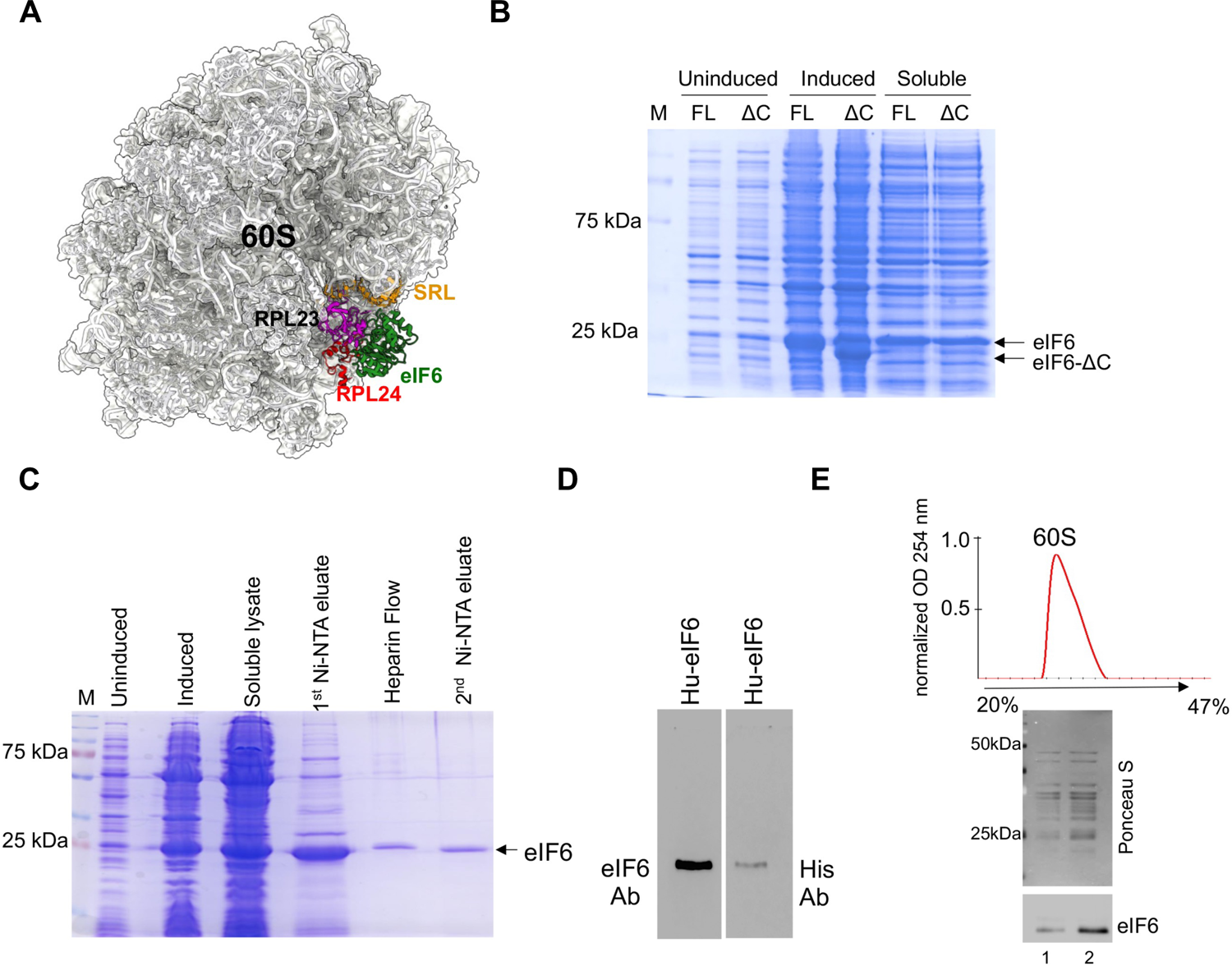
Purification of recombinant human eIF6. A) Cryo-EM structure of human eIF6 bound to 60S (PBD code:5AN9). eIF6 (green), RPL23 (uL14) (magenta), RPL24 (red) and sarcin-ricin loop (SRL) (orange) are highlighted. B) Representative Coomassie-stained gel shows induction and solubility of full-length (FL) and C-terminal deletion mutant of eIF6 (eIF6-ΔC). C) Representative Coomassie-stained gel shows eIF6 protein at various stages of protein purification process using affinity chromatography. D) Western blot analysis of purified eIF6. Recombinantly purified human eIF6 have intact N- and C-termini as protein is detected by anti-His antibody (Santa Cruz Biotechnology) targeted to the N-terminus and anti-eIF6 antibody (Cell Signaling) targeted to the C-terminus. E) Ribosome profile (top) shows 60S peak and the western blot (below) shows proteins extracted from the 60S fractions. Data is representative of three independent replicates. Lanes 1 and 2 depict two different concentrations of 60S fraction. Blots were stained with Ponceau S to detect 60S ribosomal proteins and probed with anti-eIF6 antibody to determine co-elution of eIF6 with the 60S fraction.

The C-tail of eIF6 is highly conserved in higher eukaryotes. Several phosphoproteomic studies including our study captured multisite phosphorylation of the C-tail especially at Ser235, Ser239 and Ser243 sites^9, 20, 37–42^. We found that eIF6 is critical for cells to adapt to starvation, which is regulated by phosphorylation of the C-tail^37^. This role of eIF6 is akin to the ribosomal silencing factor-RsfS (RsfA) in bacterial cells that binds to the same uL14 (rplN) interface in the large 50S subunit and is critical for cells to limit global translation to adapt to nutrient-deprivation^43^. Intriguingly, our HDX-MS analyses captured dynamic changes in the C-terminus of eIF6 upon binding to RPL23 despite the absence of direct contacts between the C-terminus of eIF6 and RPL23 in structural studies. Previously, phosphorylation of the Ser235 site was proposed to release eIF6 from the 60S^9^. However, the mechanism for this allosteric mode of regulation has remained elusive. Our CD studies show that the addition of negative charges at conserved sites of phosphorylation in the C-tail markedly influence eIF6 conformation and could explain the molecular basis for allosteric regulation of eIF6-60S interactions by the C-tail.

## Results

### Purification of recombinant human eIF6

Milligram quantities of eIF6 are required to perform detailed biophysical and structural characterization. Therefore, we first optimized the purification methodology to enhance the yield of recombinant full-length human eIF6. Previous studies have indicated that the overproduction of recombinant human eIF6 in *E. coli* is limited due to poor solubility (2 to 5%) and results in low μg/mL yield of human eIF6^36, 44^. Similarly, purification and yield of full-length *S. cerevisiae* Tif6 from *E. coli* is hindered by the disordered nature of last 20 amino acid residues in the C-terminus^34^. Removal of the last 20 residues enhanced the stability and yield of Tif6^34^. Therefore, we tested if deletion of the last 20 amino acid residues in the C-terminus of human eIF6 had a similar effect on protein yield. Deletion of the C-tail did not enhance the stability or solubility of eIF6 (Fig. 1B). This behavior is distinct from yeast Tif6 and suggests that structural contributions of the C-terminal 20 residues in eIF6 may diverge between lower and higher eukaryotes.

Co-expression of a cocktail of three (3-Ch: GroES, GroEL, TF) or five (5-Ch: DnaK-DnaJ-GrpE, GroES, GroEL) bacterial chaperones improved the solubility of eIF6 (Fig. S1A). However, the gains from this strategy to improve yields of human eIF6 were nullified by the additional purification steps required to eliminate multiple chaperones (Fig. S1A). Furthermore, co-expression of the C-terminal deletion mutant of human eIF6 along with the 5 chaperones did not further improve the solubility of eIF6 (Fig. S1B). Thus, to better refine this strategy, we systematically identified that co-expression of just the TF chaperone was sufficient to enhance solubility of human eIF6 (Fig. S1A). We next optimized the purification method for human eIF6 using affinity chromatography (Fig. 1C). Interestingly, an array of proteins coelute with human eIF6 that could potentially be ribosomal factors. Subsequent fractionation of this complex using a Heparin column results in separation of human eIF6 from all other impurities and yields ≥98% pure full-length human eIF6 with high concentration of up to 5mg/mL (Fig. 1C). Using this methodology, we also obtained high yield of yeast Tif6 without the need to delete the terminal 20 residues (Fig. S1C). The identity of full-length eIF6 was further confirmed with an anti-His-antibody that recognizes the N-terminus of eIF6 and an anti-eIF6 antibody that recognizes the C-tail of eIF6 (Fig. 1D). These results were further confirmed by mass spectrometry (data not shown). The recombinantly purified human eIF6 is active as it interacts with 60S in sucrose density gradient fractionation analysis (Fig. 1E). We further confirmed the anti-association activity of eIF6 using the subunit joining assay. We verified the purity of the isolated human 60S and 40S subunits by probing for ribosomal factors that were specific to each subunit (Fig. S1D). Negative EM images showed that 60S and 40S subunits predominantly remain dissociated at lower Mg^2+^ concentrations (Fig. 2A) and associate to form 80S only in the presence of higher Mg^2+^ concentrations (Fig. 2B). However, the association of subunits is inhibited by pre-incubation of 60S with eIF6 (Fig. 2C), which further indicates that the purified eIF6 is active. Successful overproduction and purification of milligram quantities of active human eIF6 enabled us to investigate the binding and conformational dynamics in the absence or presence of RPL23.

**Figure 2.**
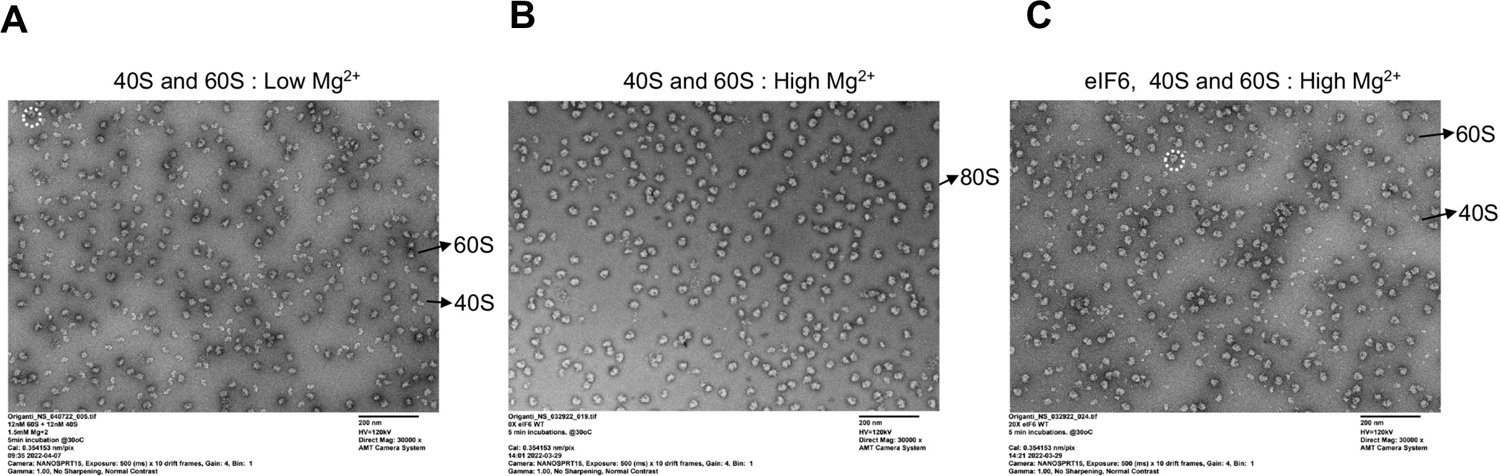
Subunit anti-association activity of recombinant human eIF6. A) Negative EM images depict 60S and 40S subunits incubated in low Mg^2+^ buffer. B) Negative EM images depict the association of 60S and 40S subunits incubated in high Mg^2+^ buffer. C) Negative EM images depict 60S and 40S subunits incubated with eIF6 in high Mg^2+^ buffer. Dotted white circles indicate the presence of a small fraction of 80S in A and C images.

### Residues in interface 1 contribute differentially to RPL23 (uL14)-eIF6 interaction

Structural studies indicate that eIF6 interacts with 60S through direct contacts with RPL23 (uL14) and is positioned proximal to RPL24 and the sarcin-ricin loop (SRL) (Fig. 1A). Majority of these contacts are between eIF6 and a short helix at the C-terminus of RPL23 (Fig. 3A). These contacts are highly conserved and observed in 60S-eIF6 structures in other organisms. However, it has never been tested if the C-terminus of RPL23 is sufficient for interaction with eIF6. Furthermore, the role of the individual contacts in promoting 60S binding to eIF6 are also unknown. This knowledge is required to probe this binding interface as a potential target for developing small molecule inhibitors against eIF6. In the human eIF6-60S structure, four residues in RPL23 (Asp127, Arg131, Asn135, and Ser138) make key contacts with eIF6 (Fig. 3B). Arg131 forms a network of side chain interactions with Asp12 and Ser190. Ser138 makes backbone interactions within RPL23 (Fig. 3B). Asp127 and Asn135 interact with Arg57 and Tyr151 of eIF6, respectively (Fig. 3B). However, in the cryo-EM structures that depict the nucleoplasmic-state A and cytoplasmic-state B of pre-60S subunits, the contacts between Tyr151 and RPL23 are varied, which indicates the dynamic nature of these interactions (Fig. S2A and B). Therefore, to delineate the contributions of specific residues in the C-terminus of RPL23 towards complex formation between RPL23 and eIF6, we performed binding experiments using SPR.

**Figure 3.**
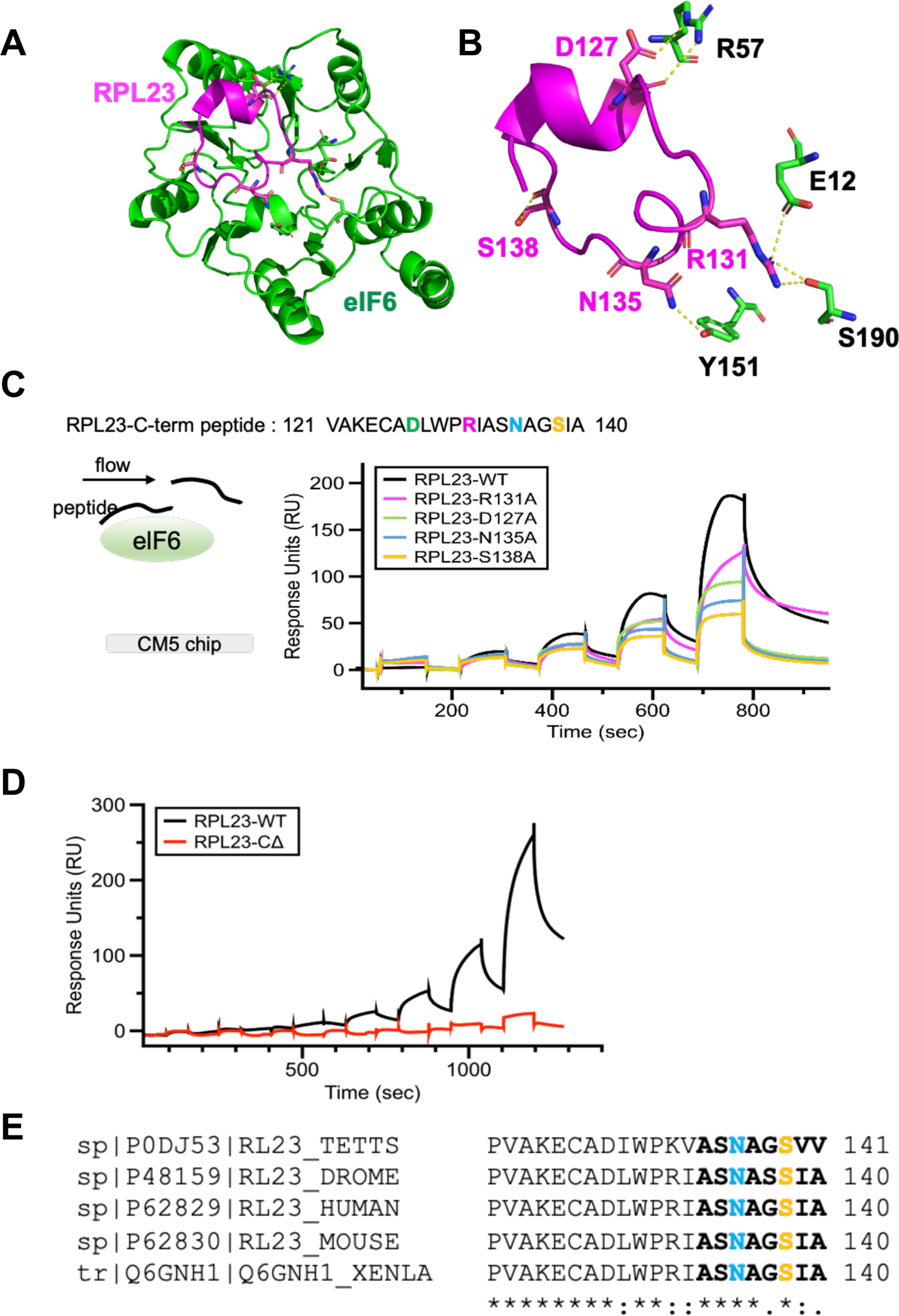
Residues in interface 1 contribute differentially to interactions between eIF6 and RPL23. A) and B) Structures highlight the location of interface 1 and the contacts between eIF6 and RPL23(uL14) (PDB code: 6UL8). C) Surface plasmon resonance experiments were performed by attaching eIF6 onto the CM5 chip and sequentially injecting increasing concentrations of RPL23 peptide. Proportional binding and dissociation are observed as a function of peptide concentration. Experiments were performed with peptides carrying various mutations as denoted. The mutations result in loss of interaction of eIF6 to varying degrees, with the most severe loss of binding observed for the Ser138-Ala substitution. D) In SPR analysis, deletion of the last eight amino acids in the RPL23 peptide results in complete loss of eIF6 binding. E) Analysis of the RPL23 sequences (last 21 residues) from various eukarya show a high degree of conservation in the C-terminus of RPL23. The terminal 8 residues in RPL23 (bold) and the Asn135 (blue) and Ser138 (orange) residues are highlighted in the sequence. Sequences were aligned using Clustal Omega. Asterisk (*), Colon (:) and dot (.) indicate identical residues, conserved and semi-conserved residues respectively.

For SPR studies, full-length elF6 was immobilized to the surface, and the C-terminal peptide of RPL23 (aa 121-140) was used in the fluid phase (Fig. 3C). We found that the RPL23 peptide bound to immobilized eIF6 in a dose-dependent manner (Fig. 3C). Association and dissociation profiles did not obey a 1:1 binding model suggesting a complex mechanism of interaction rather than non-specific binding of the peptide to the surface. The contribution of non-specific binding of the peptide to the surface was subtracted for each sensorgram. Importantly, alanine substitutions of key residues in the RPL23 peptide Arg131, Asp127 and especially, Asn135 and Ser138 showed a significant reduction (∼50% reduction at 0.5 mM) in eIF6 binding (Fig. 3C). It further highlighted the significance of the Asn135 and Ser138 for interaction with eIF6. Interestingly, Asn135 of RPL23 interacts with Tyr151 in eIF6. Suppressor mutations in Tyr151 were previously identified to rescue the growth defect of *sdo1*Δ (SBDS homolog) and *efl1*Δ (EFL1 homolog) yeast strains that mimic the slow growth phenotype of SDS^45^. Thus, our SPR analyses indicate that Tyr151 is a key residue that is important for human eIF6 interaction with RPL23 and explain the effect of Tyr151 mutation to rescue the eIF6-release defect seen in SBDS deficient yeast strains.

Somatic variants of eIF6 were recently identified in the hematopoietic cells of SDS patients and these variants were categorized as beneficial mutations that rescue the translational defect of SDS cells either by decreasing eIF6 levels or by disrupting the interactions between eIF6 and 60S^32, 33^. While majority of the somatic mutations identified in SDS led to a marked loss of eIF6 expression, the predominant eIF6^N106S^ variant and *de novo* eIF6^R61L^ variant were expressed at levels similar to wild type eIF6^32, 46^. However, the Asn106-Ser and Arg61-Leu mutations have been predicted by MD simulations to disrupt interactions with RPL23^32^. In the cryo-EM structures of human eIF6 bound to pre-60S subunits, Asn106 of eIF6 also interacts with the backbone at Ala133 and Ala136 in the last 8 amino acids of the RPL23 C-terminus (Fig. S2A, and B). Arg61 on the other hand, does not make direct contacts with RPL23, rather it stabilizes intra-eIF6 conformations through backbone interactions with several residues that in turn coordinate interactions of Asn106 and Tyr 151 (Fig. S2A and B). All three residues-Arg61, Asn106 and Tyr151 are highly conserved in eukaryotes (Fig. S2C). Since Asn106 and Tyr151 makes multiple contacts with the terminal 8 residues in RPL23, we aimed to determine if deletion of the terminal 8 amino acids in RPL23 (RPL23-CΔ) was sufficient for disrupting interactions with eIF6. The terminal 8 residues in the RPL23 peptide also harbors the key Ser138 and Asn135 sites that were identified to be critical in Fig. 3C. Deletion of the last 8 amino acids in RPL23 leads to a complete loss in binding to eIF6 (Fig. 3D). Not surprisingly, the terminal 8 residues in RPL23 inclusive of the Asn135 and Ser138 are highly conserved across species (Fig. 3E). These experiments have uncovered that the terminal 8 residues in RPL23 are critical for interaction and provide a minimal binding region that can be targeted for the development of specific inhibitors.

### HDX-MS reveals global changes in eIF6 induced by binding to RPL23

Structural information about 60S-eIF6 interactions is predominantly from cryo-EM studies that captured static snapshots of preformed complexes. However, real-time analysis of the changes in eIF6 structure upon binding to RPL23 are lacking. We therefore initiated chemical cross-linking (XL-MS) and HDX-MS experiments to investigate conformational changes in eIF6 induced upon binding to RPL23. The first step was to confirm the location of RPL23 binding on eIF6. In cross linking experiments, specific crosslinks are observed between the RPL23 C-terminal peptide and residues in direct interaction interface 1 of eIF6 (Fig. S3A). Thus, our in-solution experiments show the RPL23 peptide is binding at the expected interface on eIF6 as observed in structural studies. HDX-MS experiments were performed with human eIF6 in the absence and presence of the RPL23 peptide. We have good sequence coverage of eIF6 based on pepsin peptides and can track changes in HDX for 180 of the 265 amino acids (∼70%) (Fig. S3B). For many of the regions, multiple overlapping peptides were identified (Fig. S3B). Data was collected as a function of time (30 sec, 3 min, 30 min, 3 hrs and 24 hrs) and is described in terms of the kinetics and patterns of ΔHDX (net difference in deuterium uptake in the absence or presence of RPL23 peptide).

To construct a global landscape of the deuterium exchange and associated stability and dynamics of the hydrogen bonding network of eIF6 upon binding to RPL23, we calculated ΔHDX in the absence and presence of peptide. The change in uptake at two data points (30 sec and 3 hrs) was mapped on the structural model (Fig. 4A and B). Increased deuterium uptake (red) indicates regions that are less protected (more exposed) in the presence of RPL23 peptide. Conversely, change in the downward direction (blue) denotes decreased uptake (protected) upon peptide binding. ΔHDX comparison shows robust deuterium exchange in most regions of eIF6. Peptide level deuterium uptake curves are shown in Fig. 4 and Fig. S4 to S6. Most of the peptides that contain amino acids in interface 1 of eIF6 and associated with SDS-disease mutations including residues that render eIF6 unstable (Gly14, Arg96, Asp112, Val135) as well as residues that disrupt 60S interactions (Asn106 and Tyr151) have ΔHDX changes at early time points (Fig. 4C, D, E, F, H, I and J), which further highlights the critical residues that dictate RPL23 interactions. In terms of the kinetics of ΔHDX, both fast and slow exchange patterns are observed. We classify fast exchange as deuteration that has occurred during the early time points in our experiments (30 sec – 30 min). Slow exchange is deuterium uptake that occurs over longer periods (>30 min to 24 hrs). Differences in fast exchange are observed for multiple eIF6 peptides (e.g., Fig. 4C to K) indicating a change in protection and/or dynamics. Decreases in HDX that are maintained across time scales are consistent with protection and the adoption of a stable conformation upon binding of RPL23 (e.g., Fig. 4G and I).

**Figure 4.**
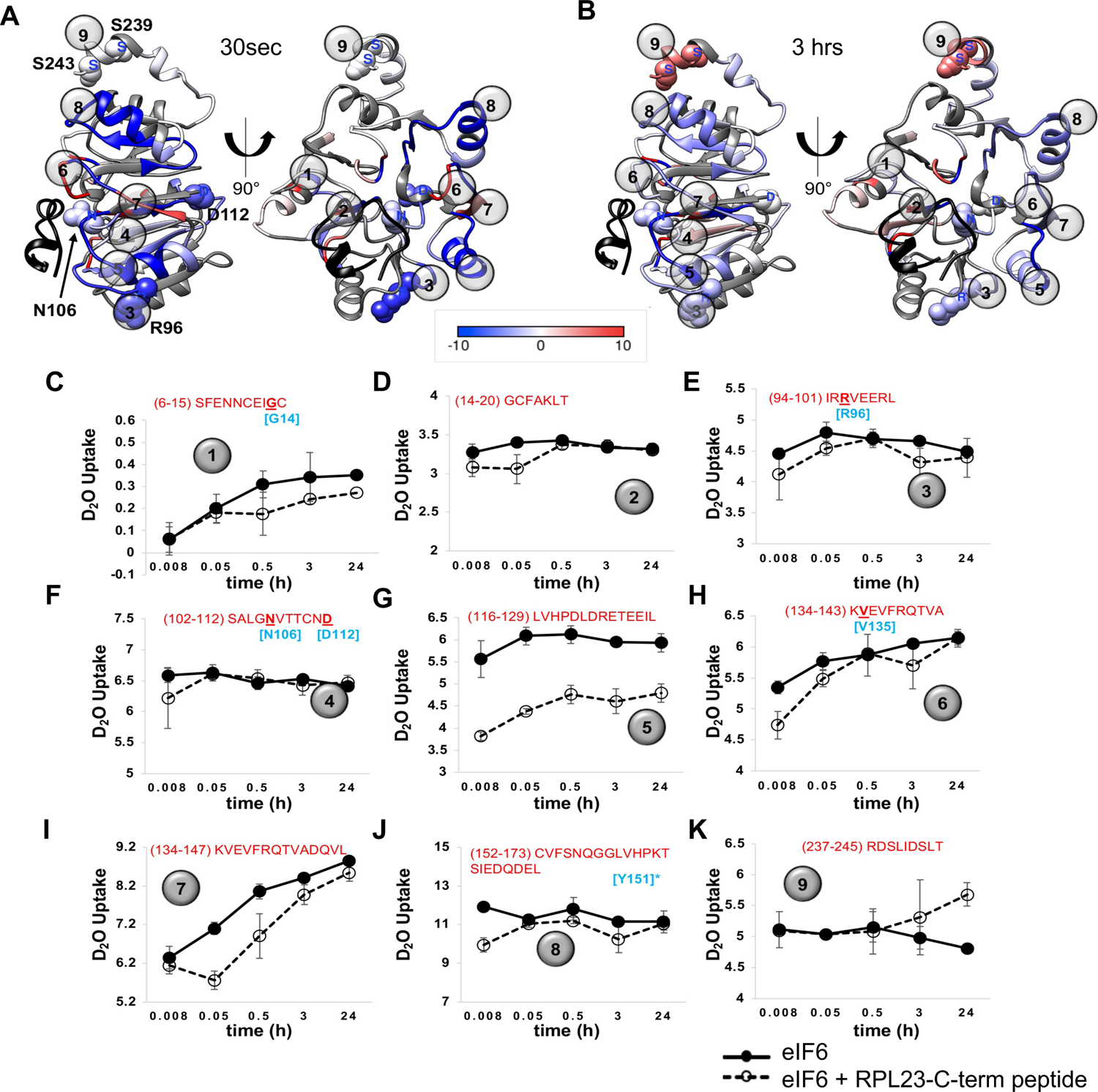
HDX-MS reveals dynamic changes in eIF6 upon RPL23 binding. A) and B) are DHDX data mapped onto the structure of eIF6 from PDB ID 6LU8. ΔHDX denotes the scale of deuterium uptake or loss measured in the absence or presence of the RPL23 peptide. The RPL23 peptide bound in the structure is shown for reference (black). Numbers 1 to 9 denote the positions of the respective peptides shown in C to K. ΔHDX changes are seen in multiple regions in eIF6 including the C-terminal helix. C to K) eIF6 peptides identified in HDX-MS analysis are shown. Data were collected as a function of time and deuterium uptake was measured in the absence of presence of the RPL23 peptide. Sequence of the individual peptides are noted in each panel. Residues noted in cyan are SDS-patient associated mutations.

Interestingly, HDX differences are also observed in the C-terminal tail of eIF6 (Fig. 4K). Several global phosphoproteomic studies have detected multiple phosphosites in the C-tail of human eIF6 especially at the Ser235, Ser239 and Ser243 residues (Fig. S7A and S7B). Two phosphosites (Ser231 and Ser233) have also been detected in the C-tail of yeast Tif6, so far (Fig. S7A and S7B)^47–52^. Phosphorylation of Ser235 in the C-terminus of eIF6 has been shown to release eIF6 from the 60S^9^. However, the mechanism of eIF6 release by the C-tail that is not in direct contact with the RPL23 interface remains elusive. The ΔHDX observed in the C-tail upon binding to RPL23 provides direct evidence for mechanical coupling between interface 1 and phosphosites on eIF6. This suggests an allosteric connection between the C-tail and interface 1.

Results show that RPL23 binding at interface 1 induces changes in stability and dynamics within eIF6 and can be better interpreted by assigning defined states. If we were to subjectively assign *state^u^*as the conformation of unbound eIF6 (in the absence of peptide) and *state^b^*as the RPL23-peptide bound, then transitions between the two states can be observed. In some regions a rapid and stable transition to *state^b^* is observed, as displayed by the constant ΔHDX over time (Fig. 4G and I). This exchange pattern is consistent with long-lasting H-bond networks and formation of stable local structures. In some cases, the transition to *state^b^* is slow (Fig. 4K and S4L) and such slow exchanges are a result of dynamic H-bond networks leading to solvent exposure and could be due to destabilization of the local structure. This infers slower remodeling rates due to conformational changes and potential allosteric processes (discussed below). Examples of such slow exchanges are observed in the disordered C-terminal region of eIF6 (Fig. 4K). In many of the regions where RPL23 peptide binding-induced changes in deuterium uptake are observed, increased protection is present (i.e., reduced deuterium uptake in the slow exchange regime, e.g., Fig. 4C, 4G and S4D). This can be interpreted as protein surfaces that become buried upon RPL23 peptide binding and are stable over the course of time (Fig 4H, 4I). If exchange is observed on the minutes to hours’ time scale, this suggests those peptide regions are transiently exposed to solvent and are therefore ‘less’ protected, hence more exposed, or dynamic (Fig. 4K).

### Interface 1 disease mutations in eIF6 alter the secondary structure

The SDS-disease mutants of eIF6 reside directly in interface 1 or close to this region (Fig. 5A and B). MD simulations have predicted that surface residues in interface 1 do not alter the overall structure of eIF6 as there is no significant effect on protein stability^32^. However, mutations in this region are expected to result in a loss of interaction with 60S. Our ability to biochemically investigate human eIF6 allow us to experimentally test whether mutations in interface 1 cause structural perturbations. The cryo-EM structures of the 60S-eIF6 complex and crystal structures of eIF6 homologs from yeast and *Tetrahymena* show eIF6 as an ordered protein. However, our HDX-MS data show extensive changes in stability and the dynamics of the hydrogen bonding network occurring within eIF6 upon RPL23 binding. Thus, in solution, the conformations of eIF6 are likely more dynamic. To gain structural insight into the solution structure of human eIF6, we performed circular dichroism (CD) analysis of the secondary structure. Simulation of the predicted CD profile for human eIF6 bound to 60S using PDBMD2CD yields a classical α-helical profile with double minima at 210 nm and 220 nm (Figure 5C). In solution CD measurements of unbound eIF6 show a slightly altered profile (Figure 5D). A strong minima is observed at ∼222 nm and reflects the α-helical content of eIF6. However, the predicted minima at ∼210 nm is overshadowed by an increase in β-sheet (and other) structural content. Analysis of the secondary structure content using BESTSEL shows ∼27% α-helix and ∼40% β-sheet (Fig. S8A). The comparison of the predicted secondary structure content from the structure of human eIF6 bound to 60S and the in-solution measurements of human eIF6 show identical α-helical content (27% each), but a considerable difference in the β-sheet content (10% versus 40%, respectively). Thus, the unbound eIF6 in solution (*state^u^*) exists in a slightly different conformation compared to when 60S bound (*state^b^*).

**Figure 5.**
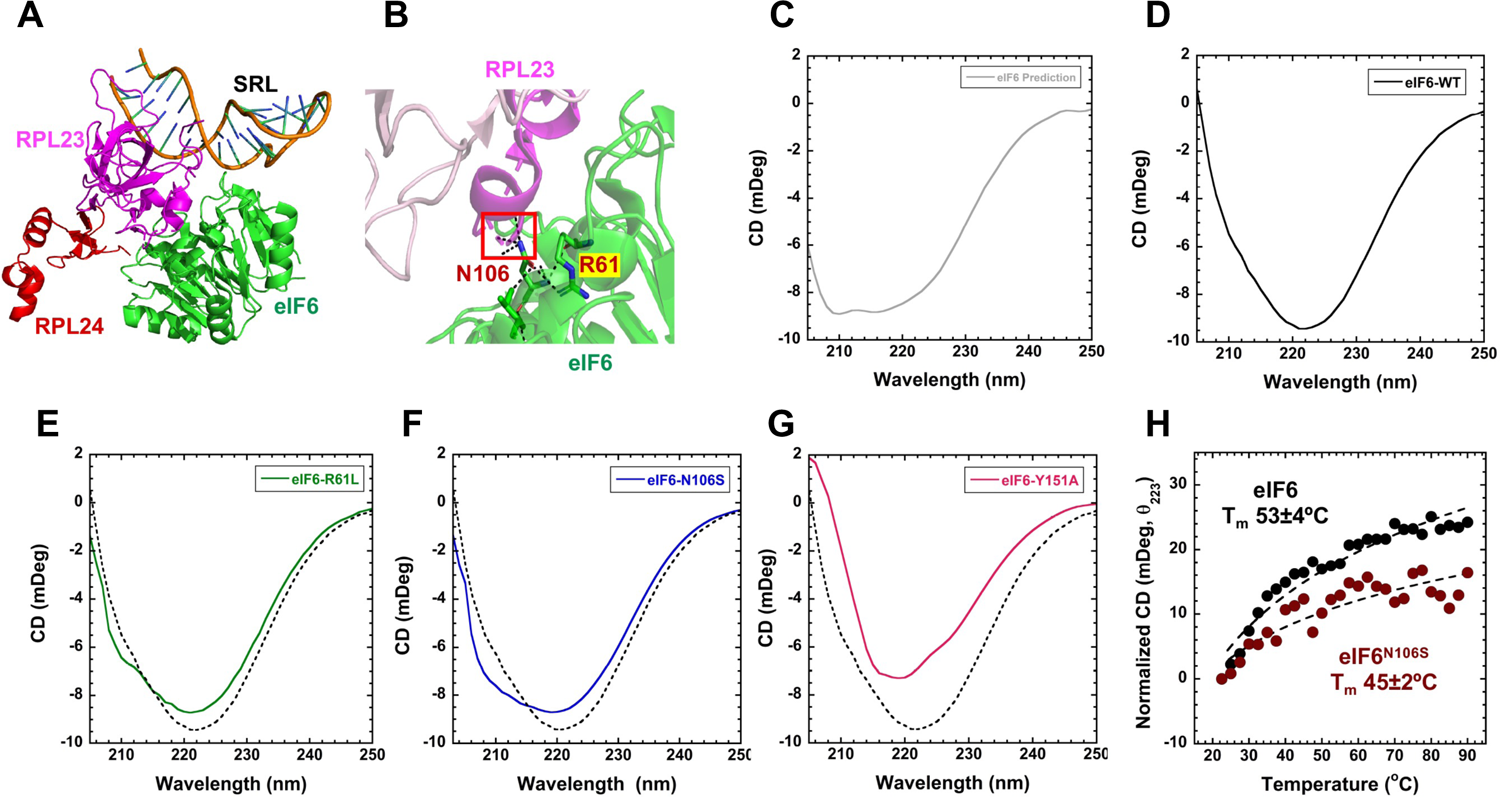
Secondary structure changes in SDS disease variants. A) Positions of RPL23 (magenta), RPL24 (red), and the SRL in relation to eIF6. B) Interactions mediated by Asn106 and Arg61 in eIF6 are depicted. C) Predicted circular dichroism (CD) spectra from the structure of human eIF6 bound to 60S (PDB code 5AN9). In solution CD measurements of D) eIF6-WT, E) eIF6^R61L^, F) eIF6^N106S^, and G) eIF6^Y151A^. Data represents three independent replicates. H) Changes in CD signal at 223 nm were recorded as a function of temperature yield T_m_ = 53±4 and 45±2 for eIF6 and eIF6^N106S^, respectively. Data is representative of three independent replicates.

Next, we measured the CD profiles for mutant eIF6 proteins that were purified similar to WT (Fig. S8B) to capture the mutation-induced changes in secondary structure. We tested the two disease-variants: eIF6^R61L^ and eIF6^N106S^. The two mutant proteins show differences in the secondary structure compared to eIF6-WT (Fig. 5D, E, F and G). eIF6^R61L^ and eIF6^N106S^ share similar α-helical content to eIF6 but show small increases in their β-sheet content. eIF6^N106^ was predicted to interact with 60S based on its position in interface 1. But MD simulations suggested that such a substitution will not alter the secondary structure of eIF6. Since *state^u^*displays a different secondary structure composition and the CD profile shows changes for the predominant eIF6^N106S^ mutation compared to wild type, we next tested the differences in their thermal stability (Fig. 5H). In contrast to computational predictions, our CD analysis show that eIF6^N106S^ is thermally less stable (Tm=45°C) compared to eIF6-WT (Tm=53°C). Thus, mutations in interface 1 also have an influence on the secondary structure of eIF6 in the unbound state.

Since Tyr151 is one of the key residues in eIF6 that interacts with Asn135 of RPL23, we also determined the effect of substituting Tyr151 with Alanine. eIF6^Y151A^ mutant shows a marked difference in CD spectra with a significant reduction of α-helical signature at 222 nm (Fig. 5G). Apart from the key interaction with Asn135 in RPL23, Tyr151 also mediates interactions with other residues within eIF6 and coordinates an extensive network of hydrogen bonding interactions that are core to the structure. Thus, Tyr151 plays an important structural role in *state^u^* of eIF6 (Fig. S9). The significance of Tyr151 was further validated in cellular studies, where we assayed the effect of homozygous expression of Tyr151Ala mutant in HCT116 cells. The Y151A mutant was expressed at slightly lower level than eIF6-WT (Fig. 6A). Expression of the eIF6 ^Y151A/Y151A^ mutant led to slower proliferation rate as shown by MTS assay (Fig. 6B). eIF6-Y151A mutation also markedly sensitized cancer cells and inhibited cell proliferation in response to nutrient stress induced by serum starvation (Fig. 6C). We did not observe any increase in cell death in mutant cells relative to wild type (Fig. 6D). This marked effect of Y151A mutation in inhibiting growth of cancer cells provides the first genetic proof of concept that targeting the eIF6-60S interaction interface could be an effective strategy for cancer therapeutics.

**Figure 6.**
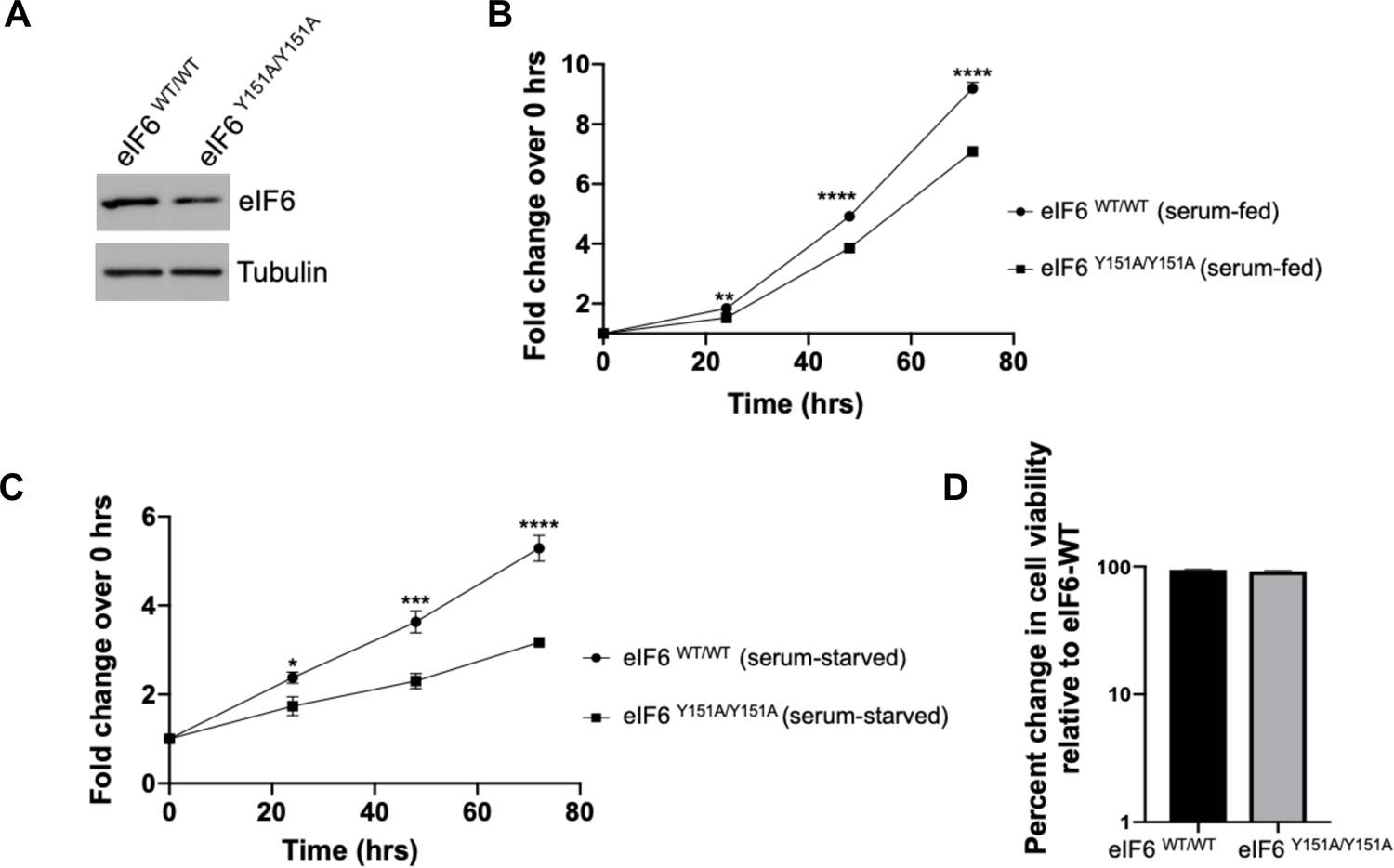
eIF6-Y151A mutation inhibits colonic cancer cell proliferation. A) Western blot shows eIF6 expression in the isogenic eIF6-WT and eIF6^Y151A/Y151A^ homozygous mutant HCT116 cells. Blots were probed with anti-eIF6 and anti-Tubulin (loading control) antibodies. Western blot is representative of three independent replicates. B) Plot depicts the fold change in cell proliferation at 24, 48 and 72 hrs relative to 0 hrs as determined by MTS assay in serum-fed cells. Values represent standard error of the mean of three independent replicates. Asterisks indicate significant differences between eIF6-WT and eIF6^Y151A/Y151A^ mutant at respective time points with p=0.0019 at 24hrs, p<0.0001 at 48 hrs and 72 hrs as determined by an unpaired two-tailed *t* test. C) Plot depicts the fold change in cell proliferation at 24, 48 and 72 hrs relative to 0 hrs as determined by MTS assay in serum-starved cells. Values indicate standard error of the mean of at least three independent replicates. Asterisks indicate significant differences between eIF6-WT and eIF6^Y151A/Y151A^ mutant at respective time points with p=0.025 at 24hrs, p=0.0012 at 48hrs and p<0.0001 at 72 hrs as determined by an unpaired two-tailed *t* test. D) Bar graph depicts percent change in cell viability of eIF6^Y151A/Y151A^ mutant relative to eIF6-WT in serum-fed cells as determined by trypan blue exclusion test.

### The C-terminus of eIF6 offers a second regulatory interface 2 for RPL23 interactions

The HDX-MS experiments uncovered allostery-driven structural changes in the C-tail of eIF6 upon binding to RPL23 in interface 1 (Fig. 4K). Several sites of phosphorylation have been identified in this region (Fig. 7A), with Ser239 and Ser243 being the predominant sites detected in the phosphoproteomic studies as indicated in Fig. S7B. Ser/Thr239 and Ser/Thr243 sites are also highly conserved in both higher and lower eukaryotes (Fig. S7A). Our previous study also detected phosphorylation of the Ser239 and Ser243 sites in serum-starved cells and showed that phosphorylation can functionally regulate eIF6^37^. In addition, our 2D-gel analysis of endogenous eIF6 in colon carcinoma cells indicates extensive phospho-modification (Fig. S10 A and B). These results are consistent with previous 2D analysis of eIF6 in malignant pleural mesothelioma cells that showed extensive phosphorylation of C-tail of eIF6 that was lost upon phosphatase treatment^20^. To better understand the effects of C-tail phosphorylation, we tested if substitution of phosphomimetic amino acids (Ser to Glu substitution) in the C-terminal region alter the secondary structure of eIF6. The phosphomimetic mutants were purified similar to WT (Fig. S10C). When negative charges are introduced in the C-terminus at the respective sites: eIF6^S239E^, eIF6^S243E^ and eIF6^S235E^ (Fig. 7B, C and D), all phosphomimetic mutants show marked changes in the secondary structures compared to WT-eIF6. Deletion of the C-terminus (eIF6^ΔC^) does not change the profile of the CD spectrum but shifts the overall profile due to the loss of the residues (Fig. 7E). As a control experiment, when Ser243 is substituted with an Ala, CD spectra are similar to WT-eIF6 (Fig. 7F) indicating that the addition of charges rather than mutation of the residue contributes to the changes in CD spectra. These CD data indicate that phosphorylation at these residues can significantly influence the overall conformation of the non-60S bound state of eIF6 (*state* ^u^). These data are in agreement with the ΔHDX changes observed in the C-terminus upon RPL23 peptide binding. It is likely that when the specific sites are phosphorylated, the C-tail binds to other regions in eIF6 and influences the secondary structure. In support of this model, the eIF6 structure from *Chaetomium thermophilum* (Tif6) shows two sulfate ions bound to the flat surface (interface 2) away from the RPL23 binding interface 1 (Fig. S11). Such intra-eIF6 interactions mediated by the negatively charged C-tail could influence the overall secondary structure and either modulate interactions with 60S or alter protein stability.

**Figure 7.**
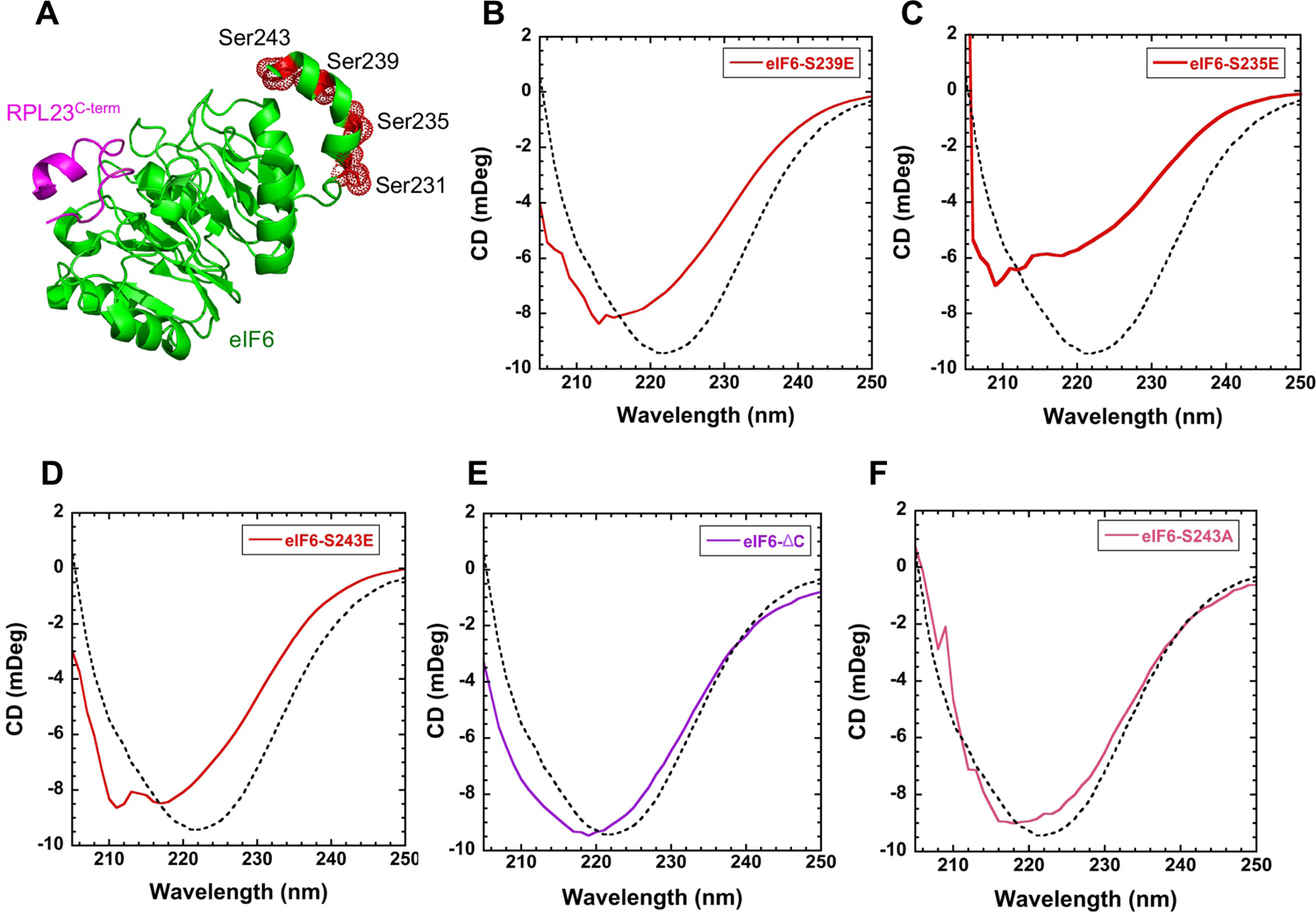
Phosphomimetic substitutions induce secondary structure changes in eIF6. A) Positions of known sites of phosphorylation in eIF6 are shown in red (PDB code 5AN9). B) In solution CD measurements of eIF6^S239E^, C) eIF6^S235E^, D) eIF6^S243E^, E) eIF6^ΔC^, and F) eIF6^S243A^. Changes in secondary structure are observed for all the phosphomimetic substitutions. eIF6^ΔC^ shows a shift in the CD spectrum due to deletion but maintains the overall profile. eIF6^S243A^ shows minimal changes in secondary structure compared to wild type eIF6. Data represents three independent replicates.

## Discussion

In this study, we show that eIF6 is conformationally dynamic and undergoes extensive conformational changes upon binding to the RPL23 interface of 60S subunit. We also establish that the terminal 8 residues in the C-terminus of RPL23 (uL14) and especially Asn135 and Ser138 residues are critical for interaction with eIF6 (Fig. 8). The positioning of these residues and their associated contacts are expected to be disrupted by the N106S mutation that is predominant in SDS patients. There is an effort to therapeutically target eIF6 by screening for compounds that broadly target eIF6 and 60S interactions^53^. In this study, by identifying the key residues of interaction, we now present a framework for a more selective target to screen for small molecule inhibitors.

**Figure 8.**
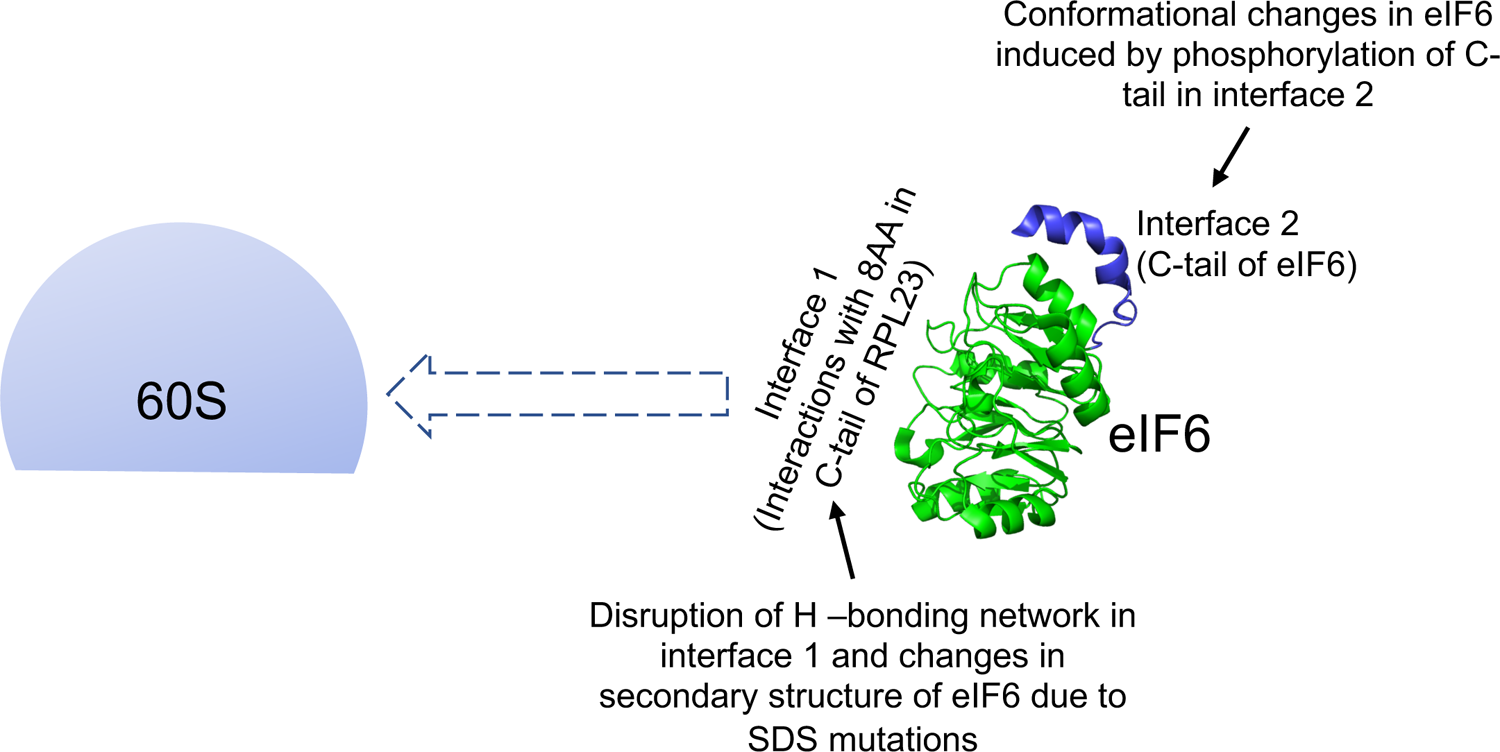
Model depicts the influence of interface 1 and 2 of eIF6 on 60S association. The key residues of interaction in interface 1 between eIF6 and the terminal 8 amino acids in the C-tail of RPL23 and conformational changes in disease variants influence the direct association of eIF6 with 60S and could influence the kinetics of eIF6 release from 60S. Phosphorylation of the C-tail of eIF6 in interface 2 and associated conformational changes of phosphomimetic mutants also influence eIF6 interactions with 60S.

In SDS hematopoietic cells, several truncating mutations of *EIF6* as well as a rare interstitial deletion in chromosome 20 that deletes *EIF6* were identified^32, 33^. In addition, several point mutants of *EIF6* were identified that were categorized as either *unstable mutants* that rendered eIF6 unstable and decreased eIF6 levels or *interaction-site mutants* that were predicted to disrupt interactions with 60S without decreasing eIF6 levels^32, 33^. Several common point mutations including the prominent R96W mutation render eIF6 unstable and these mutants are expressed poorly^32, 33^. However, the R61L mutation and the predominant N106S mutation are interaction-site mutants that are expressed at levels similar to WT but show reduced binding to the 60S subunit as shown by sucrose density gradient analysis^32^. MD simulations and homology models have predicted the effects of mutations on the overall stability and conformation of eIF6^32, 33^. For the N106S mutation, it was predicted that the sidechain of serine formed weaker H-bonds with the backbone of uL14 residues Ala133 and Ala136 or with Arg61 that weakens the H-bonding network and thereby disrupts the RPL23-interaction interface^32^. Since the simulations focused on RPL23 interactions, we wanted to determine if these mutations also affected the overall conformation of eIF6. Here, through direct testing, our CD spectra show that the secondary structure of N106S mutant is significantly altered compared to WT. This indicates that besides the influence of the side chain of the substituted Serine, the overall change in the N106S mutant conformation will further disrupt the H-bonding network in the interaction interface and weaken the binding of N106S to 60S.

Mutations of Tyr151 and Asn106 residues were identified among the gain of function alleles that suppressed the slow growth phenotype of *sdo1*Δ and *efl1*Δ yeast strains^45^. Disruption of Tyr151 residue caused a marked inhibition of cancer cell proliferation under serum-fed state. These studies indicate that targeting the interface could be an effective therapeutic strategy for cancers. Interestingly, the eIF6-Y151A caused a more profound inhibition of proliferation under conditions of nutrient deprivation. This is consistent with our previous study where we uncovered that eIF6 is important for cellular adaptation to nutrient stress and this role is conserved even in bacterial cells as shown by RsfS (RsfA) function that binds to the same uL14 (rplN) interface^37, 43^. These studies suggest that effect of eIF6 inhibitors could be enhanced by fasting. Future studies will assess the effect of targeting the eIF6-60S interaction interface *in vivo*.

Intriguingly, Tyr151 variants are yet to be identified in SDS patients, whereas N106S is a prominent mutation identified in humans and yeast. This indicates a divergence in the influence of key residues in regulating the RPL23 interface in yeast versus human. Given the marked effect of Tyr151 mutation on the secondary structure of eIF6 and cellular growth, it is possible that the clonal selection for such a mutation is limited in SDS patients. This further supports the model for a threshold of eIF6 inhibition in rescuing SDS phenotype. Somatic mutations of eIF6 in SDS patients are heterozygous. This suggests that a partial loss of eIF6 function proportional to SBDS deficiency is sufficient to rescue cellular fitness and promote clonal evolution of eIF6 mutant cells. However, any further loss of eIF6 or further decrease in association of eIF6 with 60S is likely to switch the balance to translation inhibition. A marked loss of eIF6 can lead to spurious 60S and 40S association and thereby hinder translation. It is thus possible that more severe mutations such as that of the Tyr151 residue are not as favored for clonal selection in SDS patients as they could switch the balance from rescue of cellular fitness to inhibition of growth. This threshold effect could also explain the paradox of the strategy to target the eIF6-60S interaction interface in cancers to inhibit growth whereas targeting the same interface in SDS patients can be used to rescue the slow growth phenotype and cellular fitness. In addition, normal cells that exhibit haploinsufficiency of eIF6 (eIF6^+/-^ mice) do not display changes in basal translation rates but such partial loss of eIF6 is sufficient to inhibits tumorigenesis. Thus, a dosage threshold of inhibiting eIF6 should also be taken into consideration for therapeutic screening of compounds to target eIF6-60S interactions in SBDS mutant cells versus cancers and to minimize side effects. Future studies will probe the threshold-effect of targeting eIF6 based on the cellular context and diseased state.

Almost two decades ago, phosphorylation of Ser235 was shown to release eIF6 from 60S^9^. However, the mechanism for how phosphorylation of the C-tail influences association with 60S without direct interactions with the RPL23 interface has remained unknown. Here, we show for the first time the effect of the C-terminal residues on the overall conformation of eIF6 and the dynamic solvation of the C-tail upon binding to RPL23 (Fig. 8). This suggests that the overall changes in eIF6 conformation mediated by phosphorylation of the C-tail could alter the H-bonding network at the RPL23 interface and potentially release eIF6 from 60S through allosteric regulation. However, it remains unclear as to how the C-tail contributes to the mechanism of release mediated by SBDS and EFL1 GTPase.

Since phosphorylation of endogenous eIF6 at the Ser243, Ser239 and Ser235 sites has been captured by several phosphoproteomic studies, we attempted to synthesize phospho-specific antibodies targeted against the Ser235 and Ser239 sites. However, antibody generation has been quite challenging due to the high hydrophobicity of C-tail and has yielded non-specific antibodies with enhanced cross-reactivity with the non-phosphorylated form of eIF6. Future studies will optimize antibody production to better understand the contributions of the phosphorylated form. Given the marked effect of phosphorylation on eIF6 conformation, it is likely that phosphorylation permits release of eIF6 from nascent 60S or prevents re-binding of eIF6 to recycled 60S post-termination. Studies thus far indicate that the phosphorylated fraction as well as the specific sites of phosphorylation may vary based on the physiological context and future studies will aim to better understand the mechanism and context-specific regulation of phosphorylation.

## Acknowledgements

We thank Peyton High, Yu-Fong Peng, Dr. Rahul Chadda, Dr. Nilisha Pokhrel, Dr. Elliot Corless, Dr. Jaigeeth Deveryshetty, Dr. Jinfan Wang (Stanford University), Dr. Greg Sabat (University of Wisconsin Madison-Mass Spectrometry Core), Dr. Katherine Basore and Dr. James Fitzpatrick (Washington University in St. Louis-Center for Cellular Imaging and Cryo-EM), and Genome Engineering and IPSC Center at Washington University in St. Louis for their technical support.

## Funding and Additional Information

This work was supported by grants from the National Institutes of Health R01GM143179 and R15GM126477 to S.O., R01HL150146 to N.P., R01GM130746 & R01GM133967 to E.A., and R01GM139977 & R00GM119173 to S.E.W. Funding for Proteomics, Metabolomics and Mass Spectrometry Facility at MSU was made possible in part by the MJ Murdock Charitable Trust and NIGMS of the National Institutes of Health under Award Number P20 GM103474 and *S10OD28650* to B.B.

## Data Availability

All data have been included in the manuscript.

## Materials and Methods

### Cloning and site-directed mutagenesis of eIF6

Codon-optimized full-length human *EIF6* (IDT Inc.) was cloned into MCS1 of pRSF-DUET-1 vector. Site specific mutations were introduced using Q5 -site directed mutagenesis (New England Biolabs) and primers were designed using NEBaseChanger program and were synthesized by Integrated DNA Technologies. Codon-optimized full-length *TIF6* (Genscript Inc.) was cloned into MCS1 of pRSF-DUET-1 plasmid. List of primers used are indicated in Table S1.

### Purification of recombinant full-length human eIF6

Recombinant full-length human eIF6 was co-expressed with trigger factor (TF) in *E. coli* BL21 (DE3) cells. pRSF-DUET-1-*EIF6* construct was used to express eIF6 and the pTf16 (Takara Biosciences) construct was used to express the TF chaperone. 1 to 4 L cultures were grown at 37°C in LB media supplemented with 50 μg/mL kanamycin, 34 μg/mL chloramphenicol, and 2 mg/mL L-Arabinose (to induce chaperone expression). When cultures reached OD_600_ of 0.4 to 0.5, eIF6 expression was induced with 0.4 mM IPTG and cultures were shaken for 20 hrs at 20°C. Cells were harvested by centrifugation at 4000 rpm for 20 min. For cells collected from 1L cultures, cells were resuspended in 40 mL of lysis buffer (50 mM Tris-HCl, pH 8.0, 300 mM NaCl, 10 mM imidazole, 1 mM β-mercaptoethanol, and 1 mM PMSF). In addition, 5X protease inhibitor cocktail (PIC) (Sigma-P2714), 2 μg/mL DNase, 20 μg/mL RNase and 1 mg/mL lysozyme were added to the resuspended cells. Cells were lysed by stirring for 30 min at 4°C, followed by flash freezing in liquid N_2_ and thawing in a water bath at room temperature. The freeze-thaw cycle was repeated twice, followed by Dounce homogenization at 4°C. Lysate was clarified at 17,000 rpm (rotor ID: JA 25.50) for 1 hr at 4°C. All purification steps indicated below were performed at 4°C.

The clarified lysate was applied onto a Ni^2+^-nitrolotriacetic acid (NTA) column (gravity flow) packed with 2.5 ml bed volume (BV) of agarose resin per 1 L culture. Non-specifically bound proteins were washed off with 20X bed volume of lysis buffer followed by 2 washes with 2X BV of wash buffer (50 mM Tris pH 8.0, 1 M NaCl, 60 mM imidazole, 1mM β-mercaptoethanol, 1X PMSF, and 3X PIC). Protein was eluted using 5X BV of elution buffer (50 mM Tris pH 8.0, 300 mM NaCl, 300 mM imidazole, 1 mM β-mercaptoethanol, 1X PMSF, and 5X PIC). The eluate was collected as five equal fractions and assessed by SDS-PAGE analysis. Fractions containing eIF6 were pooled, diluted 6-fold using H^0^ buffer (50 mM Tris-Cl pH 8.0 and, 1mM β-mercaptoethanol), and loaded onto to a 5 mL HiTrap-Heparin prepacked column (Cytiva Inc.) at 3 ml/min. Before loading the protein, the Heparin column was washed with 6X column volume (CV) of H^50^ buffer (50 mM Tris-Cl pH 8.0, 1 mM β-mercaptoethanol, and 50mM NaCl). eIF6 does not bind to the Heparin column under these conditions and thus flows through. eIF6 in the flow-through fraction was loaded onto a new Ni^2+^-NTA gravity flow column and washed with lysis buffer and wash buffer as described above. eIF6 was eluted using 5X BV of elution buffer (50 mM Tris pH 8.0, 300 mM NaCl, 300 mM imidazole, 1 mM β-mercaptoethanol, 1X PMSF, and 5X PIC). Five fractions of equal volume were collected and analyzed by SDS-PAGE. Fractions containing eIF6 were pooled and concentrated using an Amicon Ultra 10 kDa centrifugal concentrator and centrifuged at 4000 rpm for 10 min. To avoid mild precipitation, final volume of the concentrate from 1L culture was maintained around 500 μL. Concentrated protein was dialyzed overnight against final storage buffer (50 mM HEPES pH 7.5, 200 mM NaCl, 5 mM β-mercaptoethanol, and 10% glycerol) and briefly spun down. Small aliquots were flash frozen in liquid N_2_ and stored at −80°C. (*Note: full-length human eIF6 is prone to degradation if protein is frozen and thawed multiple times and thus, more than two freeze-thaw cycles should be avoided. After being thawed, full-length eIF6 does not tolerate re-concentration using centrifugal concentrators or overnight dialysis, at least in the buffer conditions that we tested. These processes result in either mild degradation or mild precipitation of eIF6*). Mutant eIF6 proteins were overproduced and purified using the same procedure. eIF6 concentration was measured using ε_280_ 11460 M^-1^cm^-1^. To identify the optimal co-chaperone combinations that enhance eIF6 solubility, pRSFDUET-1-*EIF6* plasmid was co-transformed with pG-KJE8 expressing five bacterial chaperones (5-Ch: DnaK-DnaJ-GrpE, GroES, GroEL) or pG-TF2 expressing 3 bacterial chaperones (3-Ch: GroES, GroEL, TF) (Takara Biosciences) in BL21 (DE3) cells. Recombinant human eIF6 were purified as described above. For these experiments, an additional purification step using size-exclusion chromatography on a Superdex S200 column (GE Healthcare) was included to remove all chaperones.

### Surface Plasmon Resonance (SPR)

SPR analyses were carried out using a BIAcore S200 instrument (GE-Healthcare). eIF6 (50 μg/mL) was solubilized in 20 mM ammonium acetate pH 4.0 and immediately immobilized on a CM5 sensor chip (4913 Response Units [RU] using NHS/EDC chemistry). After baseline stabilization, titrations were performed by injecting increasing concentrations (0-500 μM, 1:2 dilutions) of RPL23-WT (VAKECADLWPRIASNAGSIA), the site-specific mutant peptides, or the RPL23-ΔC (VAKECADLWPRI) peptide, solubilized in running buffer (20 mM HEPES pH 7.4, 150 mM NaCl, 0.01 % BSA, and 0.002% Tween 20). At the end of each titration, a solution of 20 mM HEPES pH 7.4 and 1.5 M NaCl was used to regenerate the chip. The flow rate was 25 μL/min. Peptides were solubilized in water and their concentration determined at 280 nm using the molar extinction coefficient ε_280_=5500 M^-1^ cm^-1^. To ensure reproducibility, each experiment was repeated three times. Sensograms were corrected for baseline.

### Chemical crosslinking of eIF6 to RPL23 peptide

Crosslinking reactions were performed in 50 mM HEPES pH 7.5, and 200 mM NaCl. eIF6 (24 µM final concentration) was mixed with the RPL23 peptide (0.7 mM final concentration) and crosslinker was added. 1,8-bismaleimido-diethyleneglycol (BM(PEG)2) was added to a final concentration of 3 mM and the reaction was quenched at 15 min and 1 hr with the addition of dithiothreitol (DTT, final concentration 48 mM). Quenched crosslinking reactions were then run on a Bio-Rad precast 4-20% gel (Mini-Protean TGX) at 150 V for 45 min. The gel was stained overnight with GelCode Blue Safe Protein Stain (Thermo-Fisher) and destained with deionized water. The destained gel was then subjected to a standard in gel digestion^54^ and the excised band digest was analyzed using liquid-chromatography mass spectrometry (LC-MS). LC-MS was performed on a maXis Impact UHR-QTOP instrument (Bruker Daltonics) as previously described^55^. Peptides were identified using SearchGUI v3.3.16 coupled to PeptideShaker v1.16.42 (CompOmics)^56, 57^. Crosslinked tryptic peptides were identified using the mass of the tryptic RPL23 peptide containing the crosslinkable residue as a modification and searching against the eIF6 protein sequence.

### HDX-MS of eIF6 and eIF6 bound to the RPL23 peptide

eIF6 alone (174 µM) or eIF6 (87 µM) mixed with RPL23 peptide (3.5 mM) were diluted 1:10 into deuterated buffer (50 mM HEPES, 200 mM NaCl, pD 7.5). At each time point (30 sec, 3 min, 30 min, 3 hrs, and 24 hrs), 10 µL of the HDX reaction were removed and quenched by diluting 1:6 into 0.75% formic acid (FA, Sigma), and digested for two min with porcine pepsin (0.25 mg/mL, Sigma) and vortexed every 30 seconds. Digested samples were then flash frozen and stored in liquid N_2_ until LC-MS analysis. LC-MS was carried out as described previously^58^. Before MS analysis, LC was performed on a 1290 UPLC series chromatography stack (Agilent Technologies), and peptides were separated on a reverse phase column (Phenomenex Onyx Monolithic C18 column, 100 x 2 mm) at 1 UC using a flow rate of 400 µl/min. The following conditions were used: 1.0 min, 5% B; 1.0−9.0 min, 5−45% B; 9.0−11.8 min, 45-95% B; 11.80−12.0 min, 5% B; solvent A = 0.1% FA (Sigma) in water (Thermo-Fisher) and solvent B = 0.1% FA in acetonitrile (ACN, Thermo-Fisher). MS data acquisition was carried out on a 6538 UHD Accurate-Mass QTOF LC/MS with the following settings: nebulizer set to 3.7 bar, drying gas at 8.0 L/min, drying temperature at 350 °C, and capillary voltage at 3.5 kV. Data was acquired at 2 Hz s−1 over the scan range 50−1700 m/z in positive mode. Data analysis was carried out as previously described^59, 60^ using MassHunter Qualitative Analysis (Agilent Technologies), Peptide Analysis Worksheet (PAWs, ProteoMetrics LLC), SearchGUI v3.3.16, PeptideShaker v1.16.42, HDExaminer v 2.5.1 (Sierra Analytics), and visualized using UCSF Chimera^61^.

### Western blotting

For western blot analysis, proteins were resolved by SDS-PAGE and transferred onto nitrocellulose membranes (0.45 μm; Bio-Rad Laboratories). Membranes were blocked in 5 % non-fat dry milk diluted in Tris-buffered saline with 0.1 % Tween (TBS-T) buffer. The following primary antibodies, diluted in TBS-T buffer, were used: anti-eIF6 (1:1000, overnight, D16E9, Cell Signaling), anti–His (1:1000, overnight, sc-8036, Santa Cruz Biotechnology). Anti-mouse or anti-rabbit HRP-conjugated secondary antibodies (1:30,000, 1h, Jackson Immunoresearch) were also used. For Ponceau S staining, membranes were rinsed in ultrapure water and stained with Ponceau S dye and de-stained with a brief rinse in ultrapure water. Gels and blots were imaged using iBright FL1500 imaging system (Fisher Scientific).

### Sucrose density gradient fractionation

Eukaryotic 60S (yeast) was purified as described previously^62^. 40 pmoles of eIF6 was mixed with 50 pmoles of 60S in final volume of 10 to 20 μL resuspension buffer (20 mM HEPES pH 7.3, 100 mM KCl, 5 % glycerol, 1 mM DTT, and 1 mM MgCl_2_). Reaction mixtures were incubated at 30 °C for 5 min. Samples were loaded onto 20% to 47% sucrose gradient prepared as described before^37, 63^. Gradients were centrifuged at 35,000 rpm for 2 hrs 40 min using a SW41Ti rotor (Beckman). Gradients were fractionated using a Brandel (UV) gradient fractionation system and absorbance as described previously^37, 63^, and absorbance was monitored at 254 nm. Fractions corresponding to the 60S absorbance peak were collected and subjected to trichloroacetic acid (TCA)/acetone precipitation. Precipitated proteins were analyzed by western blotting as described above.

### Purification of human 60S and 40S ribosomal subunits

60S and 40S ribosomal subunits were purified from adherent HeLa cells as described previously with a few modifications^64, 65^. HeLa cells (ATCC) were maintained in DMEM medium (Gibco) supplemented with 10% fetal bovine serum (FBS), and 100 units/mL penicillin and 100 μg/mL streptomycin (Gibco). Cells were grown to 70% to 80% confluency in 15 to 20 of 150 mm dishes. Cells were washed twice in ice-cold Phosphate Buffered Saline (PBS). Plates were briefly tilted to collect residual PBS, which was removed by aspiration. All further steps described below were carried out using RNase free tips and tubes, and buffers were made in nuclease-free ultra-pure water. Cells were scraped in lysis buffer (15 mM Tris-Cl pH 7.5, 300 mM NaCl, 7.5 mM MgCl_2_, 2 mM DTT, 1% Triton-X 100, 1 mg/mL heparin, and 0.5X cOmplete Mini EDTA-free protease inhibitor cocktail [Roche]). Lysates were rocked at 4°C for 10 min followed by centrifugation at 10,000 rpm at 4°C for 10 min. The supernatant (∼4.5 mL) was layered onto 4 mL of ice-cold 30% sucrose cushion made just before use (20 mM Tris-Cl pH 7.5, 500 mM KCl, 30% w/v RNase free sucrose, 2mM DTT, 10 mM MgCl_2_, and 0.1 mM EDTA pH 8.0) in centrifuge tubes (Beckman Coulter -catalog# 355630). Ribosomes were pelleted by centrifugation at 63,000*g* for 16–19 hrs at 4°C using a Type 80 Ti rotor (Beckman Coulter). For all ultracentrifugation steps indicated here, acceleration was set at Max and deceleration was set at Coast (no brake) setting (Beckman Coulter-Optima XPN 100 ultracentrifuge). Ribosome pellets were resuspended in 750 μL of resuspension buffer (20 mM Tris-Cl pH 7.5, 500 mM KCl, 7.5% w/v RNAse free sucrose, 2 mM MgCl_2_, 75 mM NH_4_Cl, 2 mM puromycin [Fisher Scientific], and 2 mM DTT) by gentle pipetting. The sticky pellet was gently scraped with a pipette tip, followed by gentle pipetting for 10min on ice to completely dissolve the pellet and till no chunks were visible. Ribosome pellets were then incubated at 4°C for 1 hr to facilitate 60S and 40S subunit separation. Solution was mixed by gentle pipetting and then incubated at 37°C for 1 hr. The solution was layered onto a linear 10%–30% sucrose gradient (20 mM Tris-Cl pH 7.5, 500 mM KCl, w/v RNase-free sucrose and 6 mM MgCl_2_). Step gradients were prepared by freezing individual layers in liquid N2 in centrifuge tubes (Beckman Coulter-catalog#326823, maximum volume of 38 mL) and gradients were stored in −80°C. Prior to use, gradients were thawed overnight at 4°C. A 6 to 7 mm space from the top of the tube to the sucrose layer was ensured and 750 μL of the resuspended solution was layered drop by drop onto the gradient. Gradients were centrifuged at 49,100*g* for 16 to 17 hrs at 4°C in an SW32Ti rotor (Beckman). Gradients were fractionated using a Brandel (UV) gradient fractionation system and 750 μL fractions were collected at a flow rate of 1.5 mL/min. The 40S fractions (∼4 mL total) and 60S fractions (∼7.5 mL total) were pelleted by centrifugation at 63,000*g* for 20 hrs at 4°C in a Type 80 Ti rotor (Beckman Coulter) as described above. The subunit pellets were resuspended in storage buffer (30 mM HEPES pH 7.3, 100 mM KCl, 2.5 mM Mg (OAc)_2_, 2 mM DTT and 6% w/v RNase-free sucrose). Subunits were diluted 1:500 or 1:1000 in nuclease-free ultrapure water and OD 260nm was measured (Agilent Cary 60 UV/VIS spectrophotometer). Subunit concentrations were calculated as described previously such that 1 A260 unit corresponds to 25 pmol of 60S and 50 pmol of 40S respectively^64, 65^. Subunits were aliquoted on ice in RNase-free tubes and frozen in liquid N_2_ and stored at −80°C. The 60S and 40S subunits were assessed by Coomassie staining and by western blotting using anti-RPL23 (Bethyl) and anti-RPS10 (Santa Cruz Biotechnology) antibodies diluted in 1X TBS-T buffer (1:1000, overnight) as described above.

### Secondary structure determination using circular dichroism

CD measurements were performed using a Chirascan V100 spectrometer (Applied Photophysics Inc.) A nitrogen fused set up with a cell path of 10 mm was used to perform the experiments at 20 °C. All CD traces were obtained between 200-260 nm, and background corrected using filtered CD reaction buffer (100 mM NaF, 1mM TCEP-HCl, and 5 mM Tris pH 7.5). All eIF6 proteins were first diluted to 12.5 μM in eIF6 storage buffer (50 mM HEPES pH 7.5, 200 mM NaCl, 5 mM β-mercaptoethanol, and 10% glycerol) and then diluted into 3 mL CD reaction buffer to a final concentration of 0.6 μM. 10 scans were collected and averaged per experiment using 1 nm step size and 1 nm bandwidth. Variable temperature CD was captured by monitoring the molar ellipticity at 223 nm from 20°C to 90°C in 2.5°C increments. Thermal melt data was fit to a two-state unfolding model to obtain melting temperature (Tm)^66^.

### Subunit joining assay and negative-stain electron microscopy (EM)

The reactions for subunit joining assay were set up as described previously^9^ except reactions were analyzed by negative EM. For the subunit joining assay, all reaction mixtures (50 µL) contained equal molar amounts (12 nM) of both 60S and 40S subunits in binding buffer (20 mM HEPES pH 7.3, 100 mM KCl, 1.5-10 mM Mg(OAc)_2_, 1 mM DTT, 1.5% glycerol). 20X molar excess eIF6 (280 nM) was pre-incubated with the 60S subunits for 5 min at 30°C in low (1.5 mM) Mg^2+^ binding buffer. Then, 40S was added in high Mg^2+^ binding buffer (10 mM final concentration) and incubated for 5 min at 30°C to allow for 80S formation. Reaction mixtures were immediately stained for EM imaging. For negative-stain EM, 20 µL of the subunit joining reaction mixtures were applied to plasma-cleaned carbon-coated 200 mesh copper grids (30 sec using a Gatan Solarus 950). After 1 minute incubation, the grids were washed 5X with double-stilled water and stained with 0.75% uranyl formate for 2 min. The grids were then blotted with filter paper to remove excess stain, air-dried, and imaged using a JEOL JEM-1400 120 kV transmission electron microscope (TEM) equipped with an AMT NanoSprint15 Mk-II 15-megapixel camera. EM was performed at the Washington University in St. Louis cellular imaging core.

### Cell culture

HCT116 cells (human colorectal carcinoma line) (ATCC) were cultured as described previously^37, 67^. HCT116 cells were maintained in McCoy’s 5A medium supplemented with 10% fetal bovine serum (FBS), and 100 units/mL penicillin and 100 μg/mL streptomycin (Gibco). Cells are routinely ensured to be negative for mycoplasma contamination using the mycoplasma detection kit (ATCC). For western blotting, the cells were washed twice in PBS and lysed in mammalian cell lysis buffer (MCLB) (50 mM Tris-Cl pH 8.0, 5 mM EDTA, 0.5 % Igepal, 150 mM NaCl) that was supplemented with the following inhibitors just before lysis: 1 mM phenylmethylsufonyl fluoride (PMSF), 1 mM sodium fluoride, 10 mM β-glycerophosphate, 1 mM sodium vanadate, 2 mM DTT, 1X protease inhibitor cocktail (Sigma–Aldrich), 1X phosphatase inhibitor cocktail (Santa Cruz Biotechnology). Lysates were rocked for 15 min at 4°C followed by centrifugation at 14,000 rpm for 10 min at 4°C. Blots were probed with anti-eIF6 antibody (Santa Cruz Biotechnology) (1:1000, overnight) or anti-Tubulin antibody (Cell Signaling) (1:1000, overnight) diluted in TBS-T buffer.

### Cell viability assays

5,000 HCT116 cells expressing eIF6-WT or eIF6-Y151A mutant were cultured per well of a 96-well plate in McCoy’s culture medium without phenol red supplemented with 10% FBS and 1X penicillin-streptomycin solution. For serum starvation, 7,000 HCT116 cells were cultured per well of a 96-well plate in McCoy’s culture medium supplemented with 10% FBS and after overnight incubation, cells were washed twice with 1X PBS, and cultured in McCoy’s medium supplemented with 0.1% FBS and 1X penicillin-streptomycin solution. 20 μL of MTS solution (CellTiter 96^®^ Aqueous One solution reagent, Promega) was added to each well, and the plates were incubated for 1 hr. Absorbances were read at 490 nm (Spectra Max i3x, Molecular Devices). To obtain background-corrected absorbances, average absorbance values of the control wells containing media and MTS only were subtracted from all other absorbances. Trypan blue exclusion test was performed as described before^37^.

### CRISPR/Cas9-mediated gene editing

Generation of eIF6^Y151A/Y151A^ homozygous mutant was performed at the Genome Engineering and IPSC Center (GEIC) at Washington University in St. Louis. The Y151A point mutation was introduced by nucleofecting HCT116 cells with a synthetic guide RNA (gRNA)/Cas9 ribonucleoprotein complex along with a single stranded deoxyoligonucleotides (ssODN). gRNAs were selected based on off-target analysis performed by GEIC-specific algorithms. The gRNA recognition site is 5’-AGTAGCTTCCTACTAGCACC*NGG*, with the PAM site italicized, and the ssODN has the following sequence with two phosphorothioate bonds at each end: 5’-gacagaagaaattctggcagatgtgctcaaggtggaagtcttcagacagacagtggccgaTcaggtActagtGggaagcGCTtgtgtcttcagcaatcagggagggctggtgcatcccaagacttcaattgaagaccaggatg. Both synthetic gRNA and ssODN in ultramer format were purchased from IDT. The transfected pools of HCT116 cells were analyzed by using next generation sequencing for knockin rate, and single-cell clones were obtained by sorting on a Sony sorter and screened by using next generation sequencing. Positive clones were expanded, and genotype confirmed prior to cryopreservation. All clones were negative for mycoplasma contamination and authenticated as HCT116 cells by STR profiling.

### 2D-Gel Electrophoresis

HCT116 cells were grown to 70 to 80% confluence and washed twice in PBS and collected in mammalian cell lysis buffer as described above. 2-D DIGE and Protein ID was performed by Applied Biomics, Inc (Hayward, CA). The protein extract was solubilized in 2-D lysis buffer (30 mM Tris-HCl pH 8.8, containing 7 M urea, 2 M thiourea and 4% CHAPS. Protein concentration was measured using Bio-Rad protein assay method. For Cy-Dye labeling, 30 µg of protein was mixed with 1.0 µL of diluted Cy2, and kept in dark on ice for 30 min. The labeling reaction was stopped by adding 1.0 µL of 10 mM Lysine to each sample, and incubating in dark on ice for additional 15 min. The labeled samples were then mixed 150 µg unlabeled protein. The 2X 2-D sample buffer (8 M urea, 4% CHAPS, 20 mg/mL DTT, 2% pharmalytes and trace amount of bromophenol blue), 100 μL destreak solution and Rehydration buffer (7 M urea, 2 M thiourea, 4% CHAPS, 20 mg/mL DTT, 1% pharmalytes and trace amount of bromophenol blue) were added to the labeling mix to make the total volume of 250 μL. The labeled samples are mixed then loaded into strip holder.

*IEF and SDS-PAGE:* After loading the labeled samples, IEF (pH3-10 Linear) was run following the protocol provided by GE Healthcare. Upon finishing the IEF, the IPG strips were incubated in the freshly made equilibration buffer-1 (50 mM Tris-HCl pH 8.8, containing 6 M urea, 30 % glycerol, 2 % SDS, trace amount of bromophenol blue and 10 mg/mL DTT) for 15 min with gentle shaking. The strips were subsequently rinsed in freshly made equilibration buffer-2 (50 mM Tris-HCl pH 8.8, containing 6 M urea, 30% glycerol, 2% SDS, trace amount of bromophenol blue and 45 mg/mL Iodoacetamide) for 10 min with gentle shaking. Next, the IPG strips were rinsed in the SDS-gel running buffer before transferring onto 12% SDS-gels. The SDS-PAGE were run at 15°C until the dye front bled out of the gels. Gel image was scanned immediately following the SDS-PAGE using Typhoon TRIO (GE Healthcare). The scanned images were then analyzed by Image Quant software (version 6.0, GE Healthcare). After scanning, the proteins on the gel were transferred to Immobilon PVDF membranes (EMD Millipore, Billerica, MA) at 400 mA for 2.5 hrs. Upon completion of the transfer, the membrane images were immediately scanned using Typhoon TRIO. For the western blot, membranes were blocked in 5 % BSA for 4 hrs with shaking. The membranes were then incubated overnight with shaking in primary antibody (anti-eIF6 antibody, Santa Cruz Biotechnology) in TBS-T buffer. The membranes were washed 4 times, 10 min each, with shaking in TBS-T buffer. The membranes were then incubated using Cy3 and Cy5-conjugated secondary antibody at a dilution of 1:2000 in TBS-T buffer with shaking for 2 hrs. The membranes were then washed 6 times with shaking, 10 min each, in TBST buffer. The membranes were scanned by Typhoon TRIO. The scanned images were then analyzed by Image Quant software (version 6.0, GE Healthcare).

**Main Figure Legends and Supplementary Figure Legends: Please see PPT.**

## Notes

### Competing Interest Statement

The authors have declared no competing interest.

