## Supplemental Figures for "Dynamic states of eIF6 and SDS variants modulate interactions with uL14 of the 60S ribosomal subunit"

**Figure S1.**

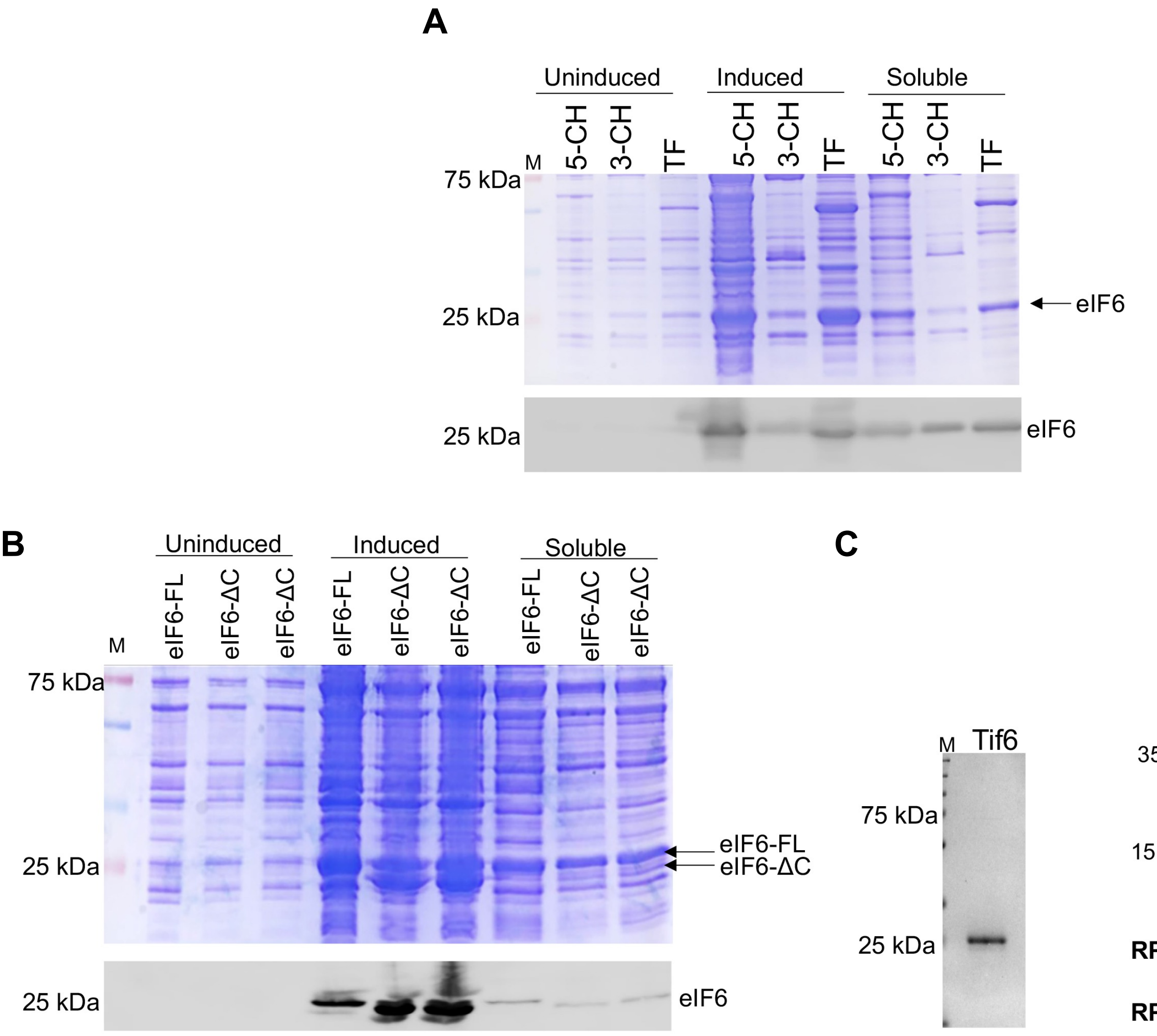

**Figure S1. Co-expression of chaperones enhance soluble production of eIF6.**
A) Representative Coomassie-stained SDS gel (top) shows the induced expression and enhanced solubility of eIF6 by co-expression of a plasmid expressing either 5 bacterial chaperones (5-CH) or 3 bacterial chaperones (3-CH) or Trigger Factor (TF) chaperone only. The corresponding western blot (bottom) was probed with anti-eIF6 antibody. B) Representative Coomassie-stained SDS gel (top) shows the expression and solubility of eIF6 by co-expression of 5-CH with either full-length (FL) eIF6 or the C-terminal deletion mutant of eIF6 (eIF6-ΔC). The corresponding western blot (bottom) was probed with anti-eIF6 antibody (Santa Cruz Biotechnology). C) Representative Coomassie-stained gel shows purified full-length yeast (*S. cerevisiae*) Tif6 protein. M-denotes molecular weight marker. D) Coomassie-stained gel (top) shows human 60S and 40S ribosomal subunits purified from HeLa cells. Purity of 60S and 40S subunits was verified by western blots (bottom) probed with anti-RPL23 and anti-RPS10 antibodies, respectively. M denotes molecular weight marker. Image is representative of at least three independent replicates.

**Figure S2.**

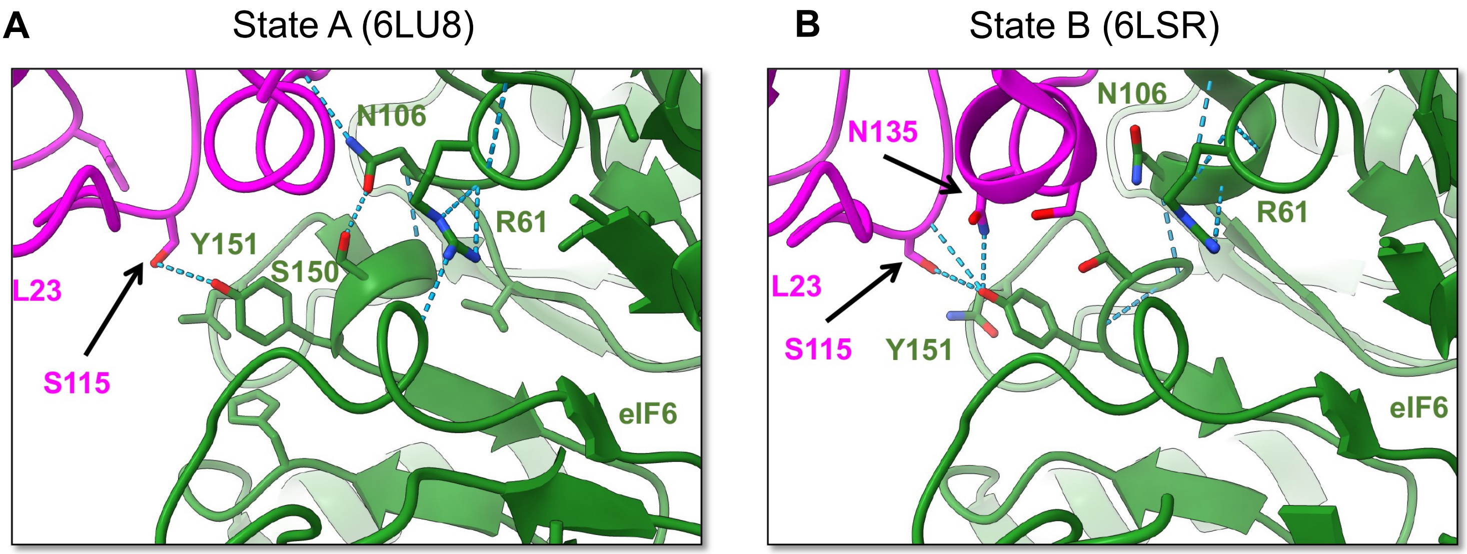

**C**

```
sp|P56537|IF6_Homo sapiens      61   RMCVGNRHGLLVPNNTTDQELQHIRNSLPDTVQIRRVEERLSALGNVTTCNDYVALVHPD   120
sp|O55135|IF6_Mus musculus      61   RMCVGNRHGLLVPNNTTDQELQHIRNSLPDSVQIRRVEERLSALGNVTTCNDYVALVHPD   120
sp|Q6ZM19|IF6_Danio rerio       61   RMCVGNRHGLLVPNNTTDQELQHIRNCLPDSVRIQRVEERLSALGNVIACNDYVALVHPD   120
sp|Q6GR45|IF6_Xenopus laevis    61   RMCVGNRHGLMVPNNTTDQELQHMRNSLPDSVRIQRVEERLSALGNVIACNDYVALVHPD   120
sp|O94476|IF6_S. pombe          61   RLTCGNRKGLLVPSSTTDNELQHLRNSLPDPVKIQRVDERLSALGNIVACNDYVALVHPD   120
sp|Q12522|IF6_S. cerevisiae     61   RMTAGNRRGLLVPTQTTDQELQHLRNSLPDSVKIQRVEERLSALGNVICCNDYVALVHPD   120
                                     *:  ***:**:**..***:****:**.*** *:*:**:********:  ***********

sp|P56537|IF6_Homo sapiens      121  LDRETEEILADVLKVEVFRQTVADQVLVGSYCVFSNQGGLVHPKTSIEDQDELSSLLQVP   180
sp|O55135|IF6_Mus musculus      121  LDRETEEILADVLKVEVFRQTVADQVLVGSYCVFSNQGGLVHPKTSIEDQDELSSLLQVP   180
sp|Q6ZM19|IF6_Danio rerio       121  LDRETEEILADTLKVEVFRQTVAEQVLVGSYCAFSNQGGLVHAKTSIEDQDELSSLLQVP   180
sp|Q6GR45|IF6_Xenopus laevis    121  LDRETEEILADVLKVEVFRQTIAEQVLVGSYCAFSNQGGLLHPKTSIEDQDELSSLLQVP   180
sp|O94476|IF6_S. pombe          121  IERETEEIIADVLDVEVFRQTVAGNVLTGSYCALSNQGALVHPRTSIQEQDELSSLLQVP   180
sp|Q12522|IF6_S. cerevisiae     121  IDRETEELISDVLGVEVFRQTISGNILVGSYCSLSNQGGLVHPQTSVQDQEELSSLLQVP   180
                                     ::*****:::*.* *******::  ::*.**** :****.*:* :**:::*:*********
```

**Figure S2. Contacts mediated by key interface residues in eIF6.** Contacts between eIF6 (green) and RPl23 (magenta) are shown in two states of human 60S maturation. A) In nucleoplasmic state A (PDB code 6LU8) and cytoplasmic state B (PDB code 6LSR), where NMD3 is stably bound, contacts between Tyr151 and RPL23 residues are varied. In both states Asn106 makes multiple contacts with the terminal 8 residues of RPL23 and Arg61 does not directly contact RPL23. E. Alignment of residues 61 to 180 in human eIF6 shows high conservation of residues mutated in SDS (shown in red): Arg61, Asn106, and Tyr151(purple) across species. Sequences were aligned using Clustal Omega. Asterisk (*), Colon (:) and dot (.) indicate identical residues, conserved and semi-conserved residues, respectively.

**Figure S3.**

**A**

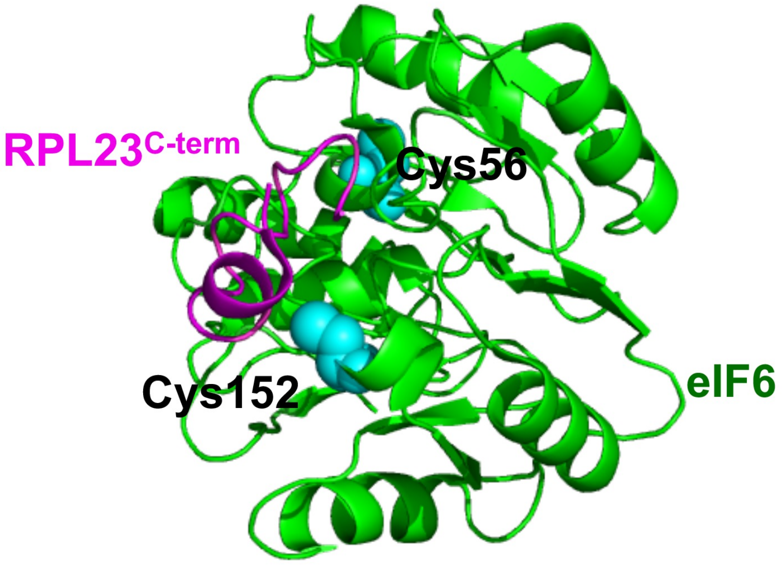

**B**

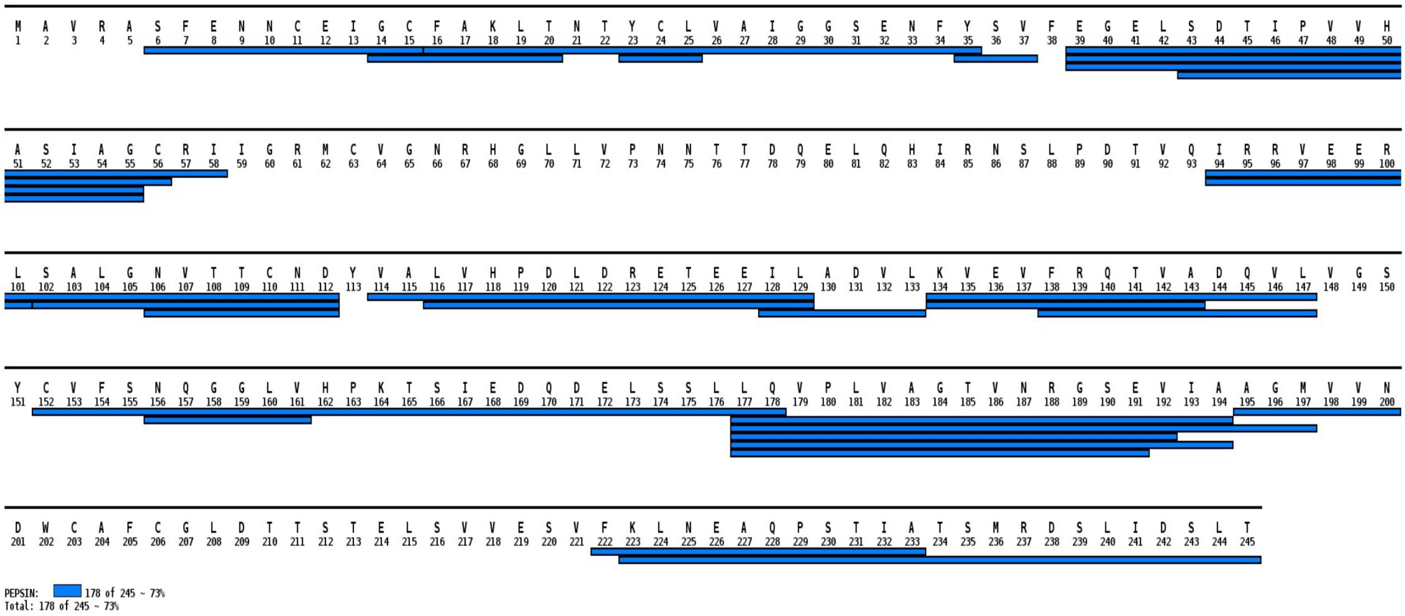

**Figure S3. HDX-MS and XL-MS analysis of eIF6 and RPL23 interactions.** A) Cys56 and Cys152 were identified in cross-linking mass spectrometry analysis of eIF6 bound to the RPL23 C-terminus peptide. The position of these residues lies in proximity to the expected site of RPL23 binding to eIF6 in the structure (PDB code 6LU8). 1,8-bismaleimido-diethyleneglycol (BM(PEG)2) was used as the crosslinker. B) Peptide coverage map of eIF6 in hydrogen-deuterium exchange mass spectrometry analysis.

**Figure S4.**

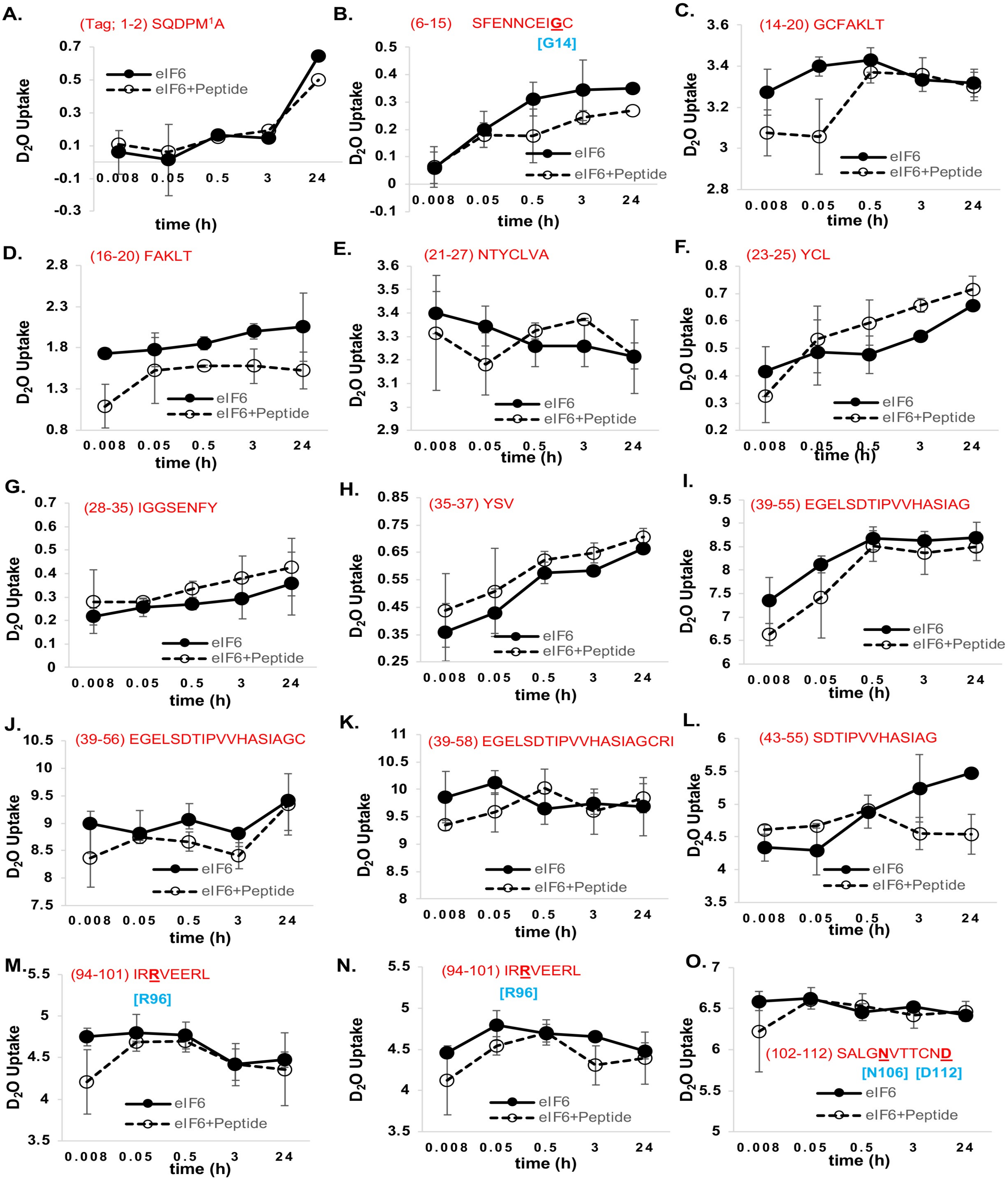

**Figure S4. Deuterium uptake in individual eIF6 peptides.** Deuterium uptake for each of the peptides are shown as a function of time. Data were collected in the absence or presence of RPL23 peptide. Residues noted in cyan were identified to be mutated in SDS patients.

#### Figure S5.

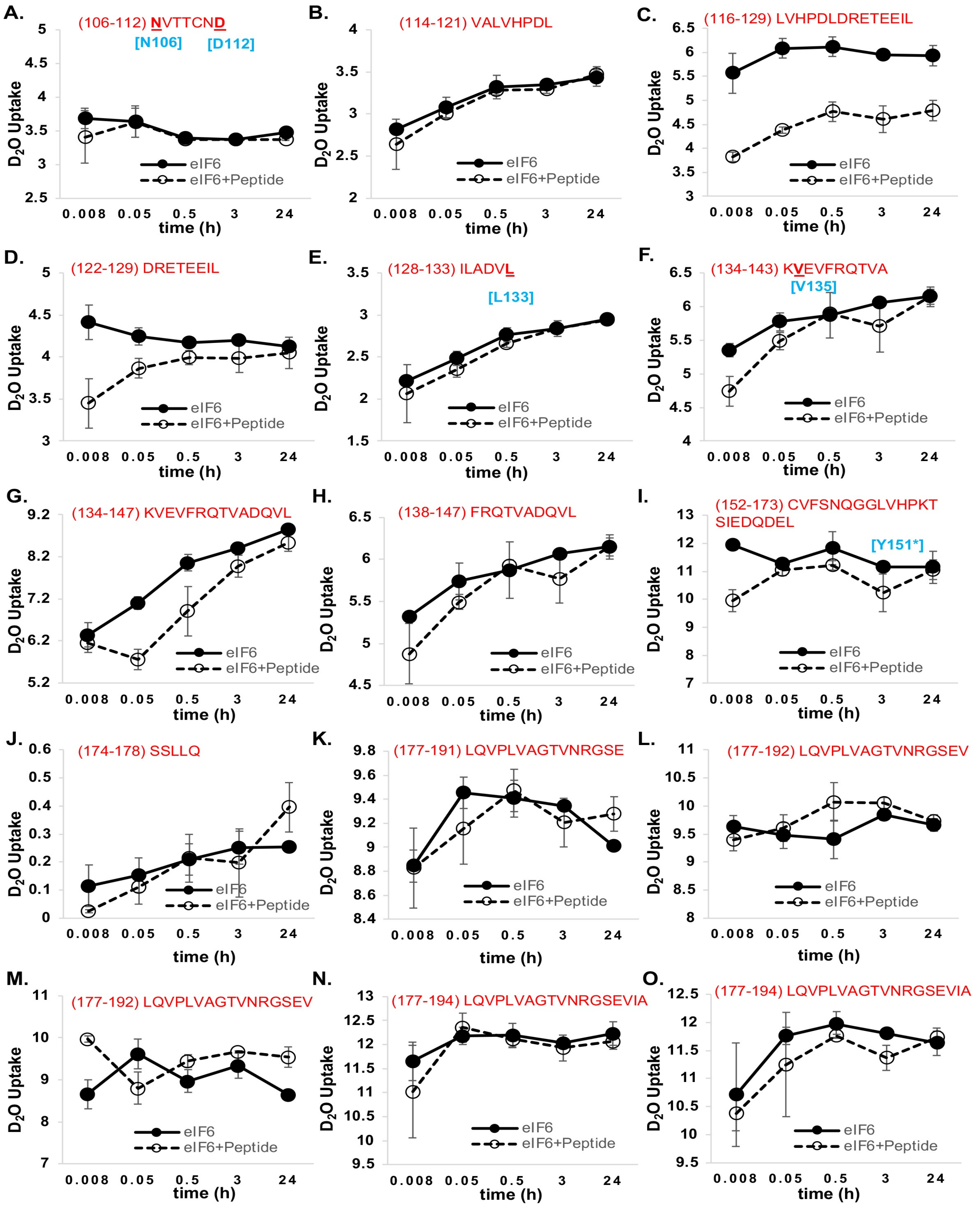

**Figure S5. Deuterium uptake in individual eIF6 peptides.** Deuterium uptake for each of the peptides are shown as a function of time. Data were collected in the absence or presence of RPL23 peptide. Residues noted in cyan were identified to be mutated in SDS patients except for residues indicated with an asterisk.

### Figure S6.

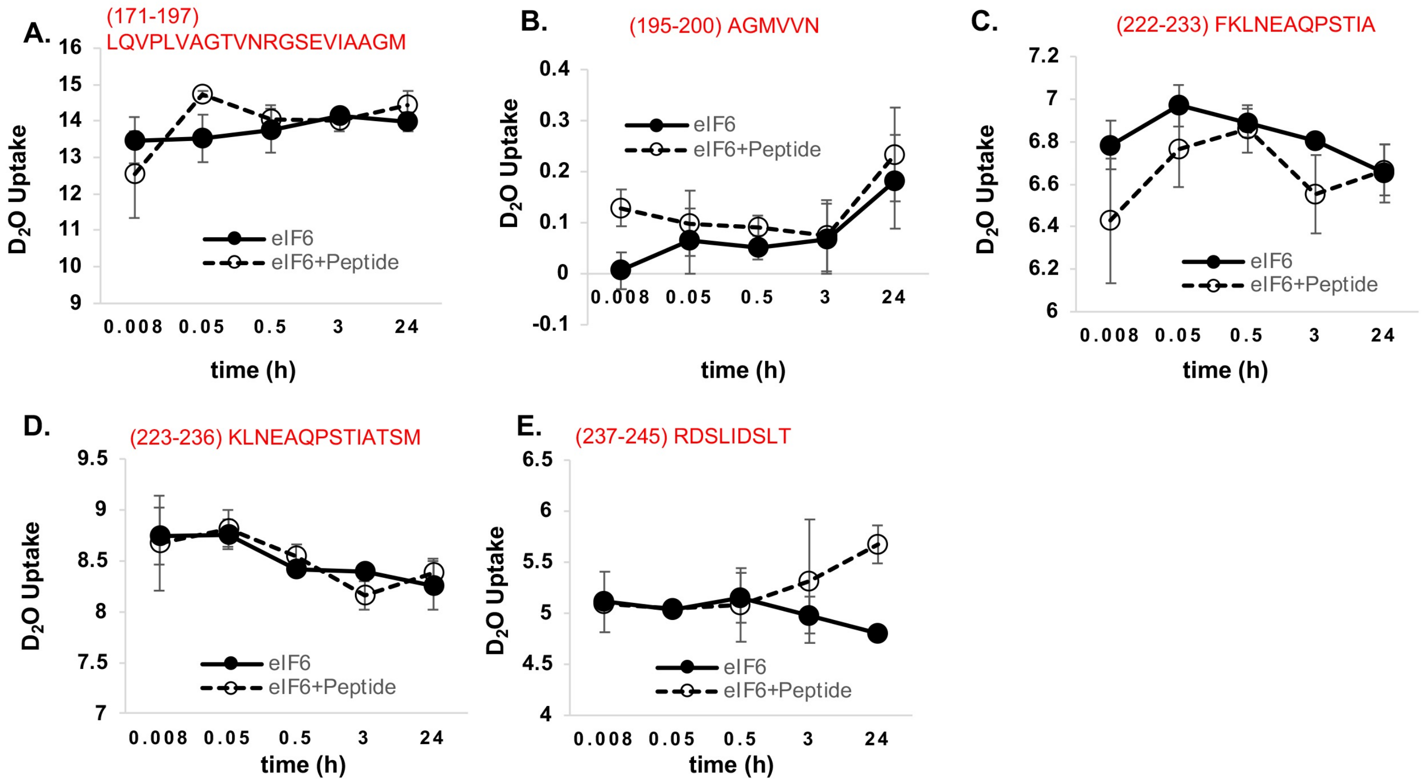

**Figure S6. Deuterium uptake in individual eIF6 peptides.** Deuterium uptake for each of the peptides are shown as a function of time. Data were collected in the absence or presence of RPL23 peptide.

### Figure S7.

**A.**

```
                                231 233 235  239  243 245
TIF6                        DAQPESISGNLRDTLIETYS 20
sp|P56537|IF6_HUMAN         EAQPSTIATSMRDSLIDSLT 20
                            :***.:*:  .:**:**:: :
```

**B.**

| Site of Phosphorylation -eIF6 | Phosphoproteomic studies that have detected phosphosites in the C-tail of human eIF6 |
|---|---|
| Ser243 | Huang H, et al. (2016) Simultaneous Enrichment of Cysteine-containing Peptides and Phosphopeptides Using a Cysteine-specific Phosphonate Adaptable Tag (CysPAT) in Combination with titanium dioxide (TiO2) Chromatography. Mol Cell Proteomics 15, 3282-3296 |
| | Mertins P, et al. (2016) Proteogenomics connects somatic mutations to signalling in breast cancer. Nature 534, 55-62 |
| | Sharma K, et al. (2014) Ultradeep human phosphoproteome reveals a distinct regulatory nature of Tyr and Ser/Thr-based signaling. Cell Rep 8, 1583-94 |
| | Mertins P, et al. (2014) Ischemia in tumors induces early and sustained phosphorylation changes in stress kinase pathways but does not affect global protein levels. Mol Cell Proteomics 13, 1690-704 |
| | Rikova K (2013) CST Curation Set: 18503; Year: 2013; Biosample/Treatment: cell line, H3122/crizotinib, geldanamycin; Disease: -; TMT: Y; Specificities of Antibodies Used to Purify Peptides prior to LCMS: pSQ, p[ST]QG Antibodies Used to Purify Peptides prior to LCMS: Phospho-ATM/ATR Substrate (S*Q) (D23H2/D69H5) Rabbit mAb Cat#: 9607 Phospho-(Ser/Thr) ATM/ATR Substrate (S*/T*QG) (P-S/T2-100) Rabbit mAb Cat#:6966 (Curated info) |
| | Schweppe DK, Rigas JR, Gerber SA (2013) Quantitative phosphoproteomic profiling of human non-small cell lung cancer tumors. J Proteomics 91, 286-96 |
| | Beli P, et al. (2012) Proteomic Investigations Reveal a Role for RNA Processing Factor THRAP3 in the DNA Damage Response. Mol Cell 46, 212-25 |
| | Kettenbach AN, et al. (2011) Quantitative phosphoproteomics identifies substrates and functional modules of aurora and polo-like kinase activities in mitotic cells. Sci Signal 4, rs5 |
| | Hsu PP, et al. (2011) The mTOR-regulated phosphoproteome reveals a mechanism of mTORC1-mediated inhibition of growth factor signaling. Science 332, 1317-22 |
| | Olsen JV, et al. (2010) Quantitative phosphoproteomics reveals widespread full phosphorylation site occupancy during mitosis. Sci Signal 3, ra3 |
| | Dephoure N, et al. (2008) A quantitative atlas of mitotic phosphorylation. Proc Natl Acad Sci U S A 105, 10762-7 |
| | Jungers CF, et al. (2020) Regulation of eukaryotic translation initiation factor 6 dynamics through multisite phosphorylation by GSK3. J Biol Chem |
| Ser239 | Mertins P, et al. (2016) Proteogenomics connects somatic mutations to signalling in breast cancer. Nature 534, 55-62 |
| | Sharma K, et al. (2014) Ultradeep human phosphoproteome reveals a distinct regulatory nature of Tyr and Ser/Thr-based signaling. Cell Rep 8, 1583-94 |
| | Imami K, et al. (2012) Temporal profiling of lapatinib-suppressed phosphorylation signals in EGFR/HER2 pathways. Mol Cell Proteomics 11, 1741-57 |
| | Klammer M, et al. (2012) Phosphosignature predicts dasatinib response in non-small cell lung cancer. Mol Cell Proteomics 11, 651-68 |
| | Kettenbach AN, et al. (2011) Quantitative phosphoproteomics identifies substrates and functional modules of aurora and polo-like kinase activities in mitotic cells. Sci Signal 4, rs5 |
| | Olsen JV, et al. (2010) Quantitative phosphoproteomics reveals widespread full phosphorylation site occupancy during mitosis. Sci Signal 3, ra3 |
| | Dephoure N, et al. (2008) A quantitative atlas of mitotic phosphorylation. Proc Natl Acad Sci U S A 105, 10762-7 |
| | Jungers CF, et al. (2020) Regulation of eukaryotic translation initiation factor 6 dynamics through multisite phosphorylation by GSK3. J Biol Chem |

**Figure S7. Key sites of phosphorylation in human eIF6 and yeast Tif6.** A) Alignment of terminal 20 residues in the C-tail of human eIF6 (P56537) and yeast TIF6 (YPR016C SGDID:S000006220) shows high conservation of Ser/Thr 239 and Ser/Thr 243 residues (in red) across species. The Ser231 and Ser233 sites (purple) in yeast Tif6 were identified as phosphosites by 2 or more phosphoproteomic studies (curated in SGD database). Sequences were aligned using Clustal Omega. Asterisk (*), Colon (:) and dot (.) indicate identical residues, conserved and semi-conserved residues, respectively. B) Table shows multiple phosphoproteomic studies that have detected endogenous phosphorylation at the highly conserved Ser239 and Ser243 sites of human eIF6 as curated in PhosphoSite Plus database.

**Figure S8.**

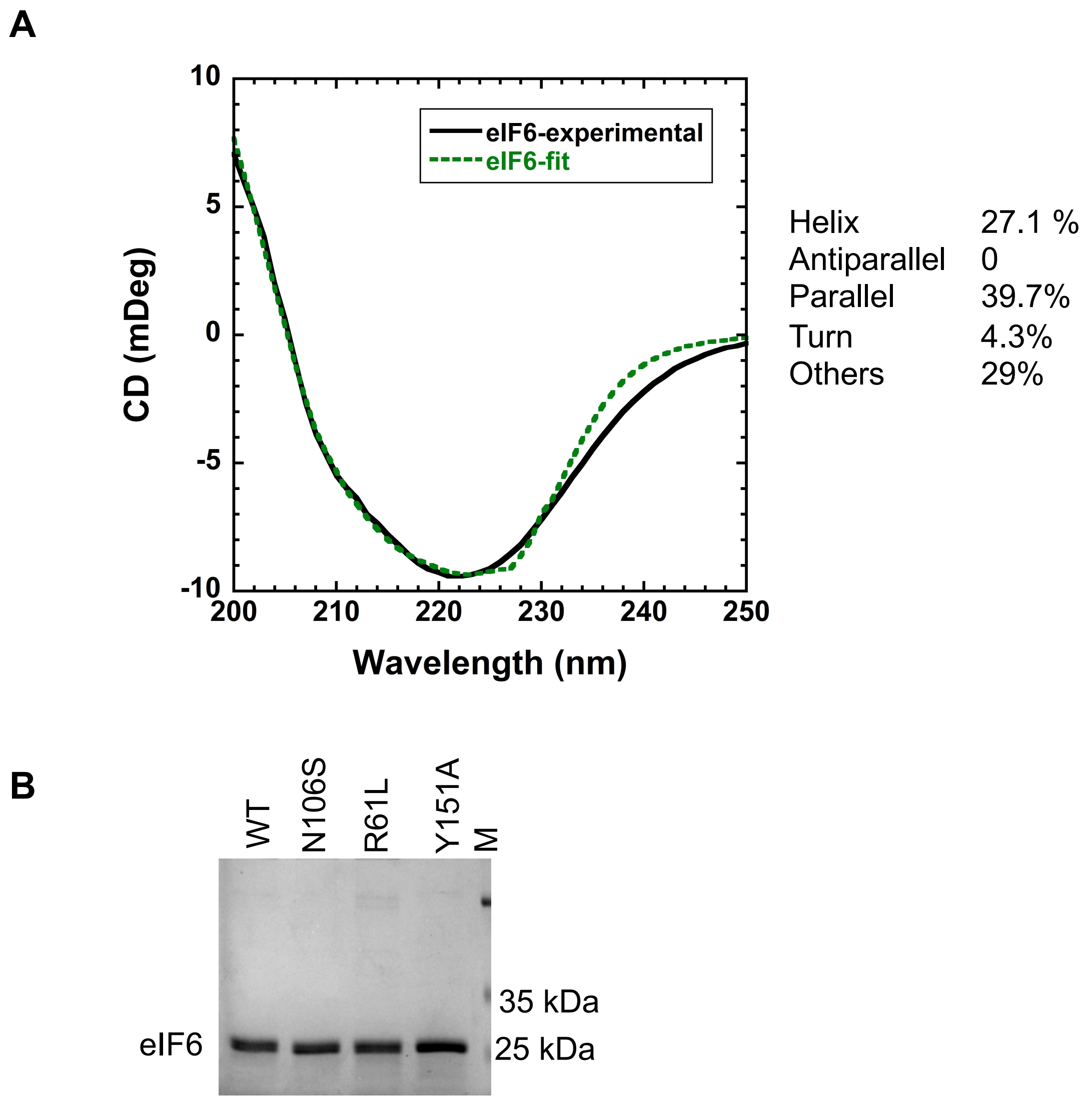

**Figure S8. Circular Dichroism (CD) analysis of eIF6 and purification of eIF6 mutants.** A) The experimental CD spectrum of eIF6 is shown (green). Data were fit with BESTSEL and the fit is shown in black. Percent distribution of secondary structure elements from the fit of the data are shown. B) Representative Coomassie-stained gel shows purified recombinant wild type (WT) and mutant eIF6 proteins. M denotes molecular weight maker. Data is representative of at least three independent replicates.

**Figure S9.**

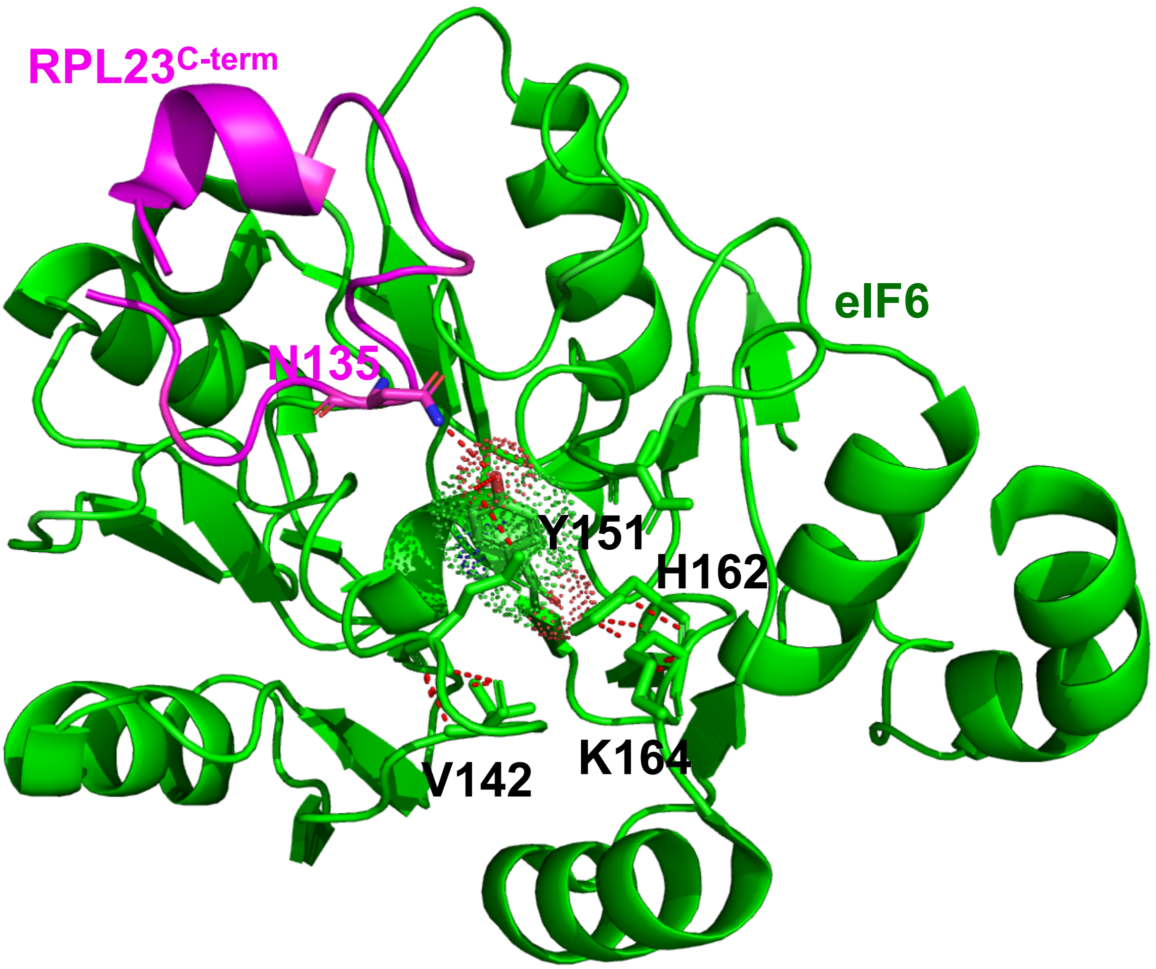

**Figure S9. Position and contacts mediated by Tyr151.** The position of Tyr151 in eIF6 is shown in red (sphere representation). Y151 contacts Asn135 in the RPL23 C-terminus and coordinates both side-chain and backbone contacts with multiple residues within the core of the eIF6 structure.

**Figure S10.**

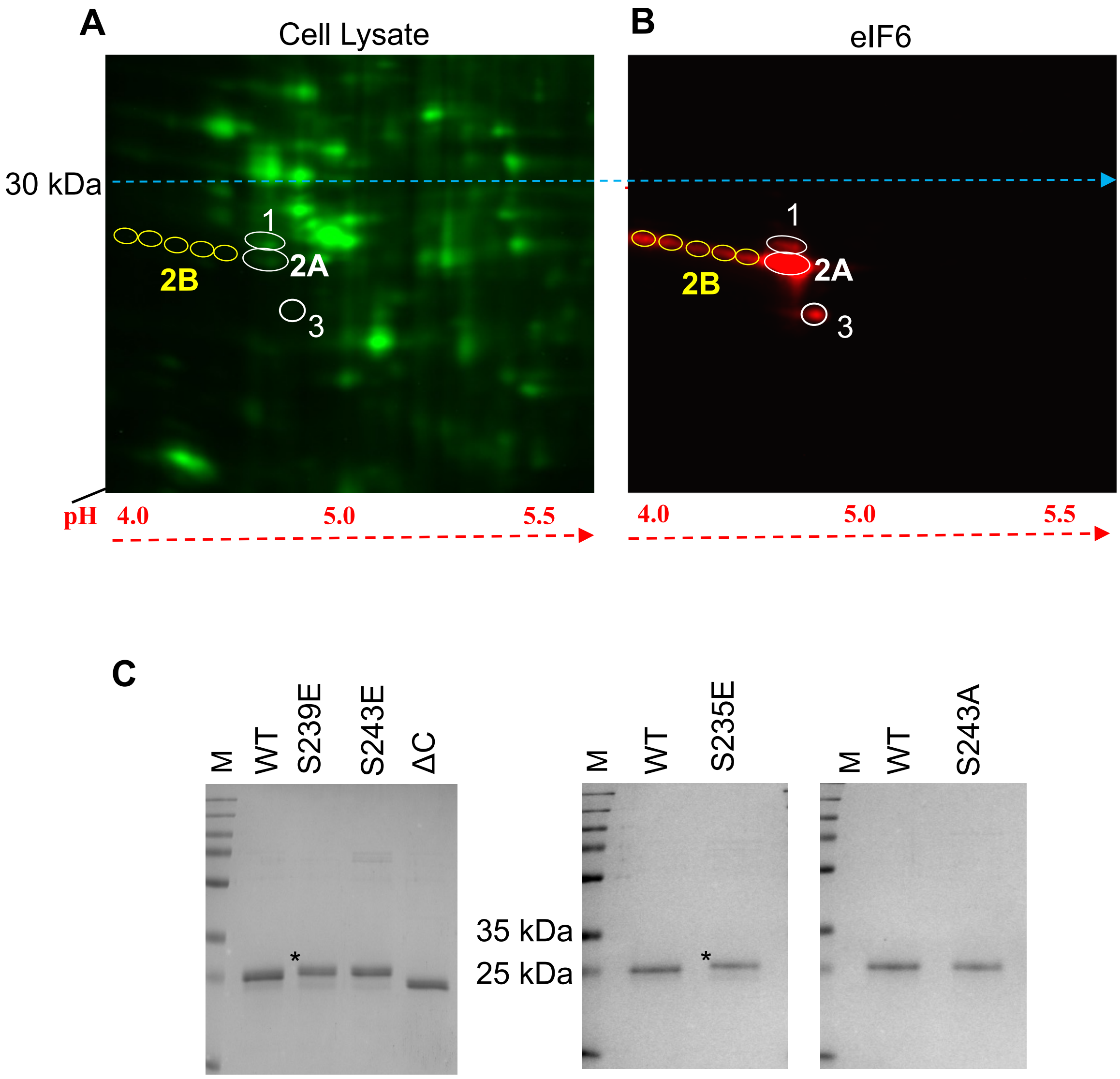

**Figure S10. 2D gel analysis of eIF6 and purification of eIF6 mutants.** A) Representative 2D-gel analysis of HCT116 cell lysate showing total protein stained with Cy2 (green). B) Representative western blot of 2D-gel probed with anti-eIF6 primary antibody (SCBT) and Cy5-conjugated anti-mouse secondary antibody. Selected spots (1, 2A, 2B and 3) depict phosphorylation of eIF6 and extracted spots were confirmed to be eIF6 by mass spectrometry. 2D-gel analysis was repeated twice. C) Representative Coomassie-stained gels show purified WT-eIF6, C-terminal deletion mutant (eIF6-ΔC), phosphomimetic mutants (S235E, S239E, S243E) and a Ser243 to Ala substitution mutant. Asterisks (*) indicate slower gel migration of the phosphomimetic mutants. Images are representative of three independent replicates. M denotes molecular weight marker.

**Figure S11.**

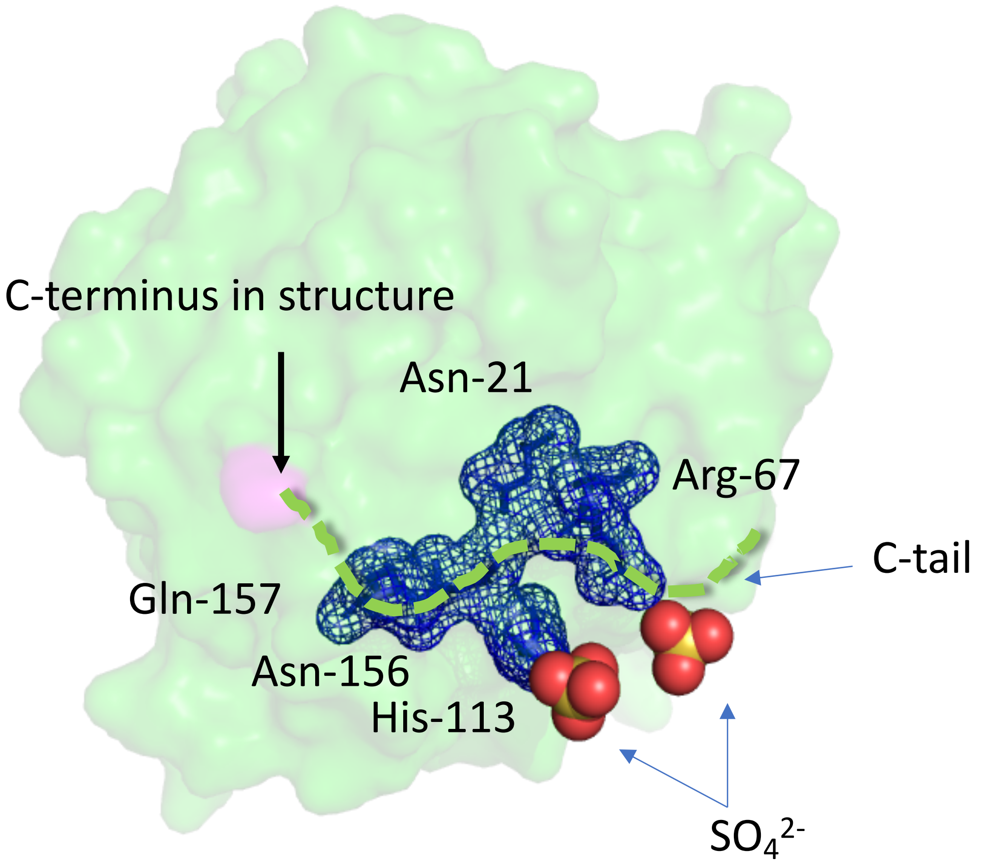

**Figure S11. Potential binding surface for the phosphorylated tail of eIF6.** Crystal structure of Tif6 from *Chaetomium thermophilum* (PDB code 5M3Q) is shown and the residues in interface 2 of Tif6 are shown in blue. The positive charges in this surface are bound to two sulfate ions in the structure. The green dotted line represents a potential binding path for the C-terminal region of eIF6 upon phosphorylation. The negative changes upon phosphorylation could be positioned similar to the sulfate ions in the structure.

**Table S1.**

| Primer | | Sequence (5' to 3') | Annealing Temp |
|---|---|---|---|
| eIF6-Y151A | Fwd | AGTGGGGTCTGCTTGCGTTTTTTC | 59°C |
| | Rev | AAGACTTGGTCAGCGACA | |
| eIF6- N106S | Fwd | GCTTTGGGGAGCGTCACCACT | 66°C |
| | Rev | AGACAAACGTTCCTCTACGC | |
| eIF6-R61L | Fwd | ATCATCGGACTCATGTGTGTCG | 63°C |
| | Rev | ACGGCAACCTGCGATAGA | |
| eIF6-ΔC | Fwd | CAAGTTAAATTAGGCTCAGCC | 56°C |
| | Rev | AATACAGATTCTACCACC | |
| eIF6-S239E | Fwd | CATGCGCGATGAACTGATTGATAG | 56°C |
| | Rev | GACGTAGCGATTGTAGAG | |
| eIF6-S243E | Fwd | CCTGATTGATGAACTGACCTGAGTCGAC | 62°C |
| | Rev | CTATCGCGCATGGACGTA | |
| eIF6-S243A | Fwd | CCTGATTGATGCTCTGACCTGAGTCGACAAG | 65°C |
| | Rev | CTATCGCGCATGGACGTA | |
| eIF6-S235E | Fwd | AATCGCTACGGAAATGCGCGATAGC | 64°C |
| | Rev | GTAGAGGGCTGAGCCTCA | |

**Table S1.** List of all forward (Fwd) and reverse (Rev) primers indicating primer names, sequences, and respective annealing temperatures used in this study.
